# Aging-associated changes in transcriptional elongation influence metazoan longevity

**DOI:** 10.1101/719864

**Authors:** Cédric Debès, Antonios Papadakis, Sebastian Grönke, Özlem Karalay, Luke Tain, Athanasia Mizi, Shuhei Nakamura, Oliver Hahn, Carina Weigelt, Natasa Josipovic, Anne Zirkel, Isabell Brusius, Konstantinos Sofiadis, Mantha Lamprousi, Yu-Xuan Lu, Wenming Huang, Reza Esmaillie, Torsten Kubacki, Martin R. Späth, Bernhard Schermer, Thomas Benzing, Roman-Ulrich Müller, Adam Antebi, Linda Partridge, Argyris Papantonis, Andreas Beyer

**Affiliations:** Cluster of Excellence Cellular Stress Responses in Aging-associated Diseases (CECAD), University of Cologne, 50931 Cologne, Germany; Max Planck Institute for Biology of Ageing, Joseph-Stelzmann-Str. 9b, 50931 Cologne, Germany; Department II of Internal Medicine and Center for Molecular Medicine Cologne, University of Cologne, Faculty of Medicine and University Hospital Cologne, Cologne, Germany; Center for Molecular Medicine Cologne, Faculty of Medicine and University Hospital of Cologne, University of Cologne, 50937 Cologne, Germany; Institute for Genetics, Faculty of Mathematics and Natural Sciences, University of Cologne, 50923 Cologne, Germany; Institute of Healthy Ageing, Department of Genetics, Evolution and Environment, UCL, London WC1E 6BT, UK; Institute of Pathology, University Medical Centre Göttingen, 37075 Göttingen, Germany

## Abstract

Physiological homeostasis becomes compromised during aging, as a result of impairment of cellular processes, including transcription and RNA splicing. However, the molecular mechanisms leading to the loss of transcriptional fidelity are so far elusive, as are ways of preventing it. Here, we profiled and analyzed genome-wide, aging-related changes in transcriptional processes across different organisms: nematode worms, fruit flies, mice, rats and humans. The average transcriptional elongation speed (Pol-II speed) increased with age in all five species. Along with these changes in elongation speed we observed changes in splicing, including a reduction of unspliced transcripts and the formation of more circular RNAs. Two lifespan-extending interventions, dietary restriction and lowered insulin/Igf signaling, both reversed most of these aging-related changes. Genetic variants in Pol-II that reduced its speed in worms and flies increased their lifespan. Similarly, reducing Pol-II speed by overexpressing histone components, to counter age-associated changes in nucleosome positioning, also extended lifespan in flies and the division potential of human cells. Our findings uncover fundamental molecular mechanisms underlying animal aging and lifespan-extending interventions, and point to possible preventative measures.

## Introduction

Aging impairs a wide range of cellular processes, many of which affect the quality and concentration of proteins. Among these, transcription is particularly important, because it is a main regulator of protein levels *(1-3)*. Transcriptional elongation is critical for proper mRNA synthesis, due to the co-transcriptional nature of pre-mRNA processing steps such as splicing, editing, and 3’ end formation *(4, 5)*. Indeed, dysregulation of transcriptional elongation results in the formation of erroneous transcripts and can lead to a number of diseases *(6,7)*. During aging, animal transcriptomes undergo extensive remodeling, with large-scale changes in the expression of transcripts involved in signaling, DNA damage responses, protein homeostasis, immune responses, and stem cell plasticity *(8)*. Furthermore, some studies uncovered an age-related increase in variability and errors in gene expression *(9-11)*. Such prior work has provided insights into how the transcriptome adapts to, and is affected by, aging-associated stress. However, it is not known if, or to what extent, the transcription process itself affects or is affected by aging.

In this study, we used high-throughput transcriptome profiling to investigate how the kinetics of transcription are affected by aging, how such changes affect mRNA biosynthesis, and to elucidate the role of these changes in age-related loss of function at the organismal level. We document an increase in Pol-II elongation speed with age across five metazoan species, a speed reduction under lifespan-extending conditions, and a causal contribution of Pol-II elongation speed to lifespan. We thus reveal an association of fine-tuning Pol-II speed with genome-wide changes in transcript structure and chromatin organization.

## Results

The translocation speed of elongating RNA polymerase II (Pol-II) can be measured using RNA sequencing (RNA-seq) coverage in introns. This is because Pol-II speed and co-transcriptional splicing are reflected in the characteristic saw-tooth pattern of read coverage, observable in total RNA-seq or nascent RNA-seq measurements *(12,13)*. Read coverage generally decreases 5’ to 3’ along an intron, and the magnitude of this decrease depends on Pol-II speed: the faster the elongation, the shallower the slope *(14-16)*. High Pol-II speeds result in fewer nascent transcripts interrupted within introns at the moment when the cells are lysed. Thus, by quantifying the gradient of read coverage along an intron, it is possible to determine Pol-II elongation speeds at individual introns (**Fig. 1a,b**). Note that this measure is only weakly associated with the expression level of the transcript (**Extended Data Table 1**) *(17)*. To monitor how the kinetics of transcription changes during aging, we quantified the distribution of intronic reads resulting from RNA-seq in five animal species: the worm *C. elegans*, the fruit fly *D. melanogaster*, the mouse *M. musculus*, the rat *R. norvegicus (18)*, and humans *H. sapiens*, at different adult ages (**Extended Data Table 2** and **Methods**), and using diverse mammalian tissues (brain, liver, kidney, whole blood), fly brains, and whole worms. Human samples originated from whole blood (healthy donors, age 21-70), and from two primary human cell lines (IMR90, HUVEC) driven into replicative senescence. After filtering, we obtained between 518 and 7994 introns that passed quality criteria for reliable Pol-II speed quantification (see **Methods**). These different numbers of usable introns mostly result from inter-species variation in intron size and number, and to some extent from variation in sequencing depth. To rate the robustness of Pol-II speed changes across biological replicates, we clustered samples based on their ‘speed signatures’, i.e. on the detected elongation speeds across all introns that could be commonly quantified across each set of experiments. We observed largely consistent co-clustering of samples from the same age across species, whereas young and old samples mostly separated from each other (**Extended Data Fig. 1**). This suggests that age-related speed changes were consistent across biological replicates and reliably quantifiable in independent measurements. We observed an increase of average Pol-II elongation speed with age in all five species and all tissue types examined (**Fig. 1c** and **Extended Data Fig. 2**). Changes in Pol-II speed did not correlate with either the length of the intron or with its position within the gene (**Extended Data Table 1**). The observed increase in Pol-II elongation speed was even more pronounced after selecting introns with consistent speed changes across all replicates (i.e., always up or down with age; **Extended Data Fig. 3**). This result is non-trivial, because our analysis also revealed introns with a consistent reduction in Pol-II speed. In order to confirm our findings with an orthogonal assay, we monitored transcription kinetics in IMR90 cells using 4sU-labelling of nascent RNA. After inhibiting transcription with 5,6-dichloro-l-β-D-ribofuranosyl benzimidazole (DRB), we conducted a pulse-chase-like experiment quantifying 4sU-labeled transcripts at four time points after transcription release (i.e., at 0, 15, 30 and 45 min). This enabled us to quantify Pol-II progression into gene bodies (see **Methods** for details) and confirmed our results based on intronic slopes using proliferating (young) and senescent (old) IMR90. Pol-II speed measurements from the 4sU-based assay showed significant correlation with those from the slope-based assay (**Fig. 1d**), with Pol-II speed increasing on average in both approaches (**Fig. 1e, Extended Data Fig. 4**). Note that, although many individual genes showed a decrease in elongation speed with aging in both assays, the majority exhibited increased speed.

**Fig. 1:**
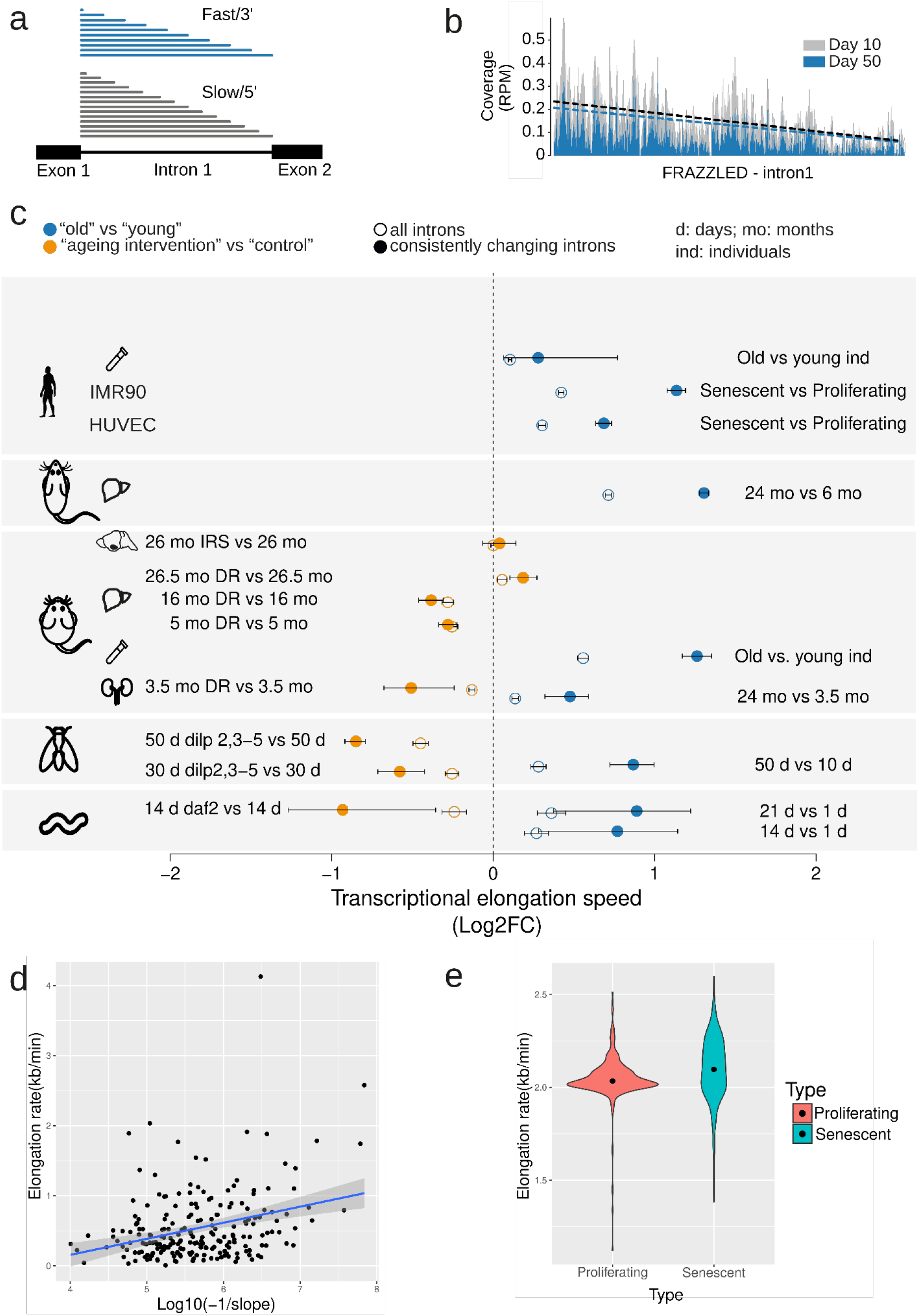
Pol-II elongation speed increases with age and is slowed-down by reduced insulin signaling and dietary restriction (DR) in multiple species. **(a)** Schematic representation of read coverage along introns in total RNA seq. Intronic reads represent transcriptional production at a given point in time. A shallower slope of the read distribution is a consequence of increased Pol-II elongation speed. **(b)** Exemplary read distribution in the FRAZZLED intron 1 with coverage in reads per million (RPM) for *D. melanogaster* at age day 10 (grey) and day 50 (blue). **(c)** Log2 fold change of average Pol-II elongation speeds in worm (whole body), fruit fly (brains), mouse (kidney, liver, hypothalamus, blood), rat (liver), human blood, HUVECs: umbilical vein endothelial cells; IMR90: fetal lung fibroblasts). Error bars show median variation ±95% confidence interval (Wilcoxon signed rank test). Empty circles indicate results using all introns passing the initial filter criteria, while full circles show results for introns with consistent effects across replicates. Number of introns considered (*n*) ranged from 518 to 7994. **(d)** Transcriptional elongation speed estimate from 4sUDRB-seq in IMR90 cells versus intronic slopes for 217 genes for which elongation speed could be estimated using both assays. Each dot represents one gene (Pearson correlation=0.313, p-value=2.5e-06). **(e)** Distributions of elongation speeds in IMR90 cells based on 4sUDRB-seq. The black dot indicates the average speed. The difference between speeds is statistically significant(paired Wilcoxon test, p-value = 2.13e-10). The same genes (464 genes) were used for both conditions (see Methods for details).

To assess whether known lifespan-extending interventions, inhibition of insulin/insulin-like growth factor signaling (IIS) and dietary restriction, affected Pol-II speeds, we sequenced RNA from IIS mutants, using *daf-2* mutant worms at day 14 and fly brains from *dilp2-3,5* mutants at day 30 and day 50, as well as hypothalamus from aged wild type and *IRS1*-null mice. We also sequenced RNA from kidney and liver of dietary restricted (DR) and *ad libitum*-fed mice. In all comparisons, except *IRS1-*null mice and livers from 26 months old DR mice, lifespan-extending interventions resulted in a significant reduction of Pol-II speed. Pol-II elongation speeds thus increased with age across a wide range of animal species and tissues, and this increase was, in most cases, reverted under lifespan-extending conditions (**Fig. 1**).

Although Pol-II speed changed consistently with age across replicates (**Extended Data Fig. 1**), we did not observe specific classes of genes to be affected across models. To determine whether genes with particular functions were more strongly affected by age-related Pol-II speed changes, we performed gene set enrichment analysis on the 200 genes with the highest increase in Pol-II speed during aging in worms, fly brains, mouse kidneys and livers, and rat livers. Only very generic functional classes, such as metabolic activity, were consistently enriched across three or more species (**Extended Data Fig. 5**). Thus, no specific cellular process appeared to be consistently affected across species and tissues. Next, we examined age-associated gene expression changes of transcription elongation regulators. We observed that some regulators (e.g., PAF1, THOC1) were consistently downregulated across species during aging (**Extended Data Fig. 6**), which was also confirmed using gene set enrichment analysis (**Extended Data Fig. 7**). These expression changes potentially represent a compensatory cellular response to a detrimental increase in transcriptional elongation speeds.

To determine if changes in Pol-II speed are causally involved in the aging process, we used genetically modified worm and fly strains carrying point mutations in a main Pol-II subunit that reduce its elongation speed (*C. elegans, ama-1* (m322) mutant *(20)*; *D. melanogaster, RpII215*^*C4*^ mutant *(21)*). We sequenced total RNA from wild type and “slow” Pol-II mutant worms (whole animal at day 14) or fly heads, at day 10 and 50. Measurements of elongation speeds confirmed the expected reduction of average Pol-II speeds in both *C. elegans ama-1* (m322) and *D. melanogaster RpII215*^*C4*^ (**Fig. 2a**). To assess whether Pol-II speed and its associated maintenance of transcriptional fidelity also affected aging of the whole organism, we measured survival of these animals. Slowing down Pol-II increased lifespan in both worms and fruit flies (median lifespan increase of ∼20 % in *C. elegans* and in ∼10 % *D. melanogaster*; **Fig. 2b** and **Extended Data Fig. 8a**). CRISPR/Cas9 engineered reversal of the Pol-II mutations in worms restored lifespan essentially to wild-type levels (**Extended Data Fig. 8b**). Furthermore, mutant worms displayed higher pharyngeal pumping rates at older age compared to wild type worms, suggesting that healthspan was also extended by slowing down Pol-II elongation speed (**Extended Data Fig. 9**).

**Fig. 2:**
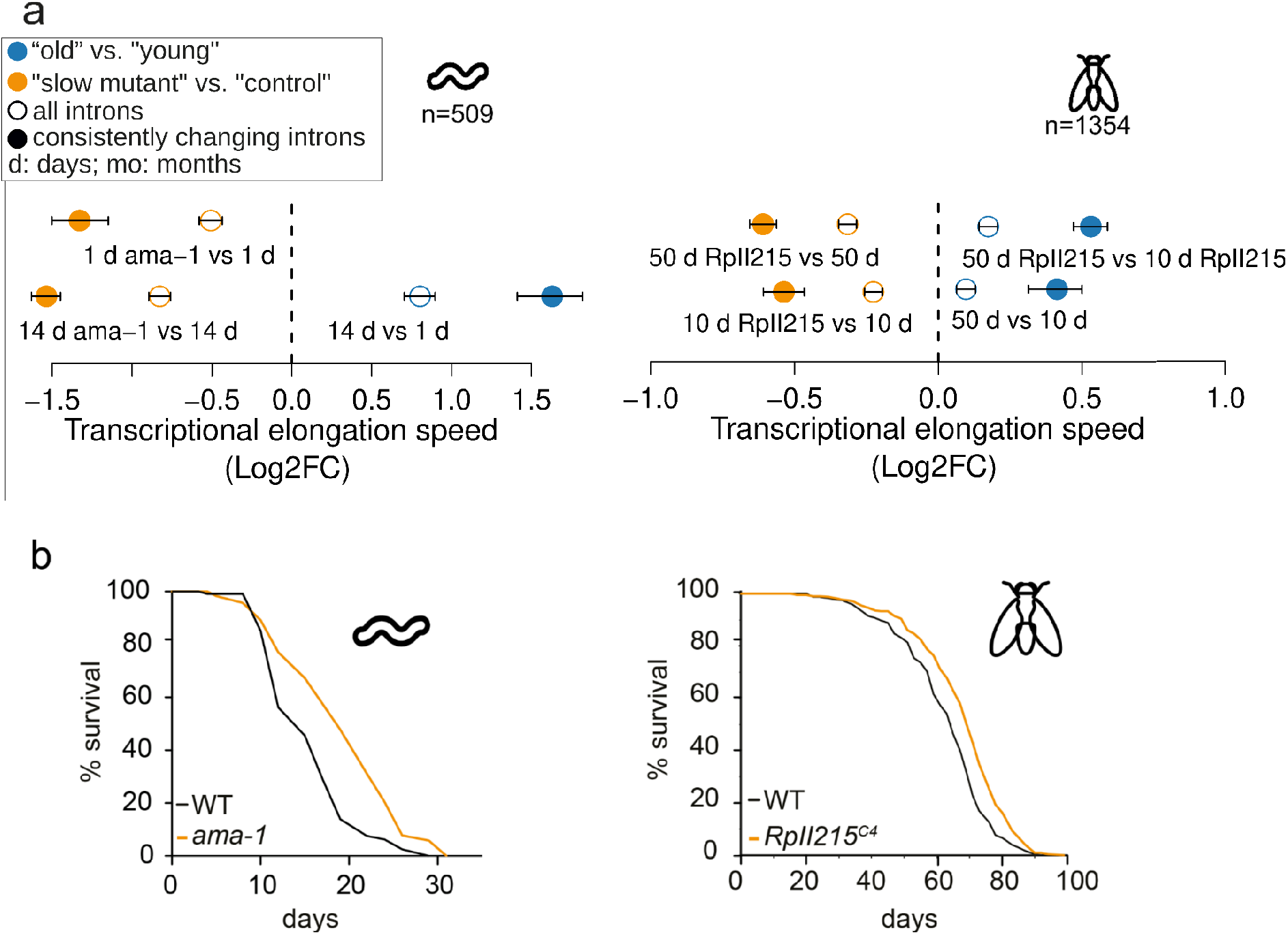
Molecular and lifespan effects of reduced Pol-II elongation speed in *C. elegans and D. melanogaster*. **(a)** Differences of average Pol-II elongation speeds between Pol-II mutant and wild-type worms (**left**) and flies (**right**), and changes of average Pol-II elongation speeds with age in flies (right). Error bars show median variation ±95% confidence intervals. All average changes of Pol-II elongation speeds are significantly different from zero (P < 0.001; paired Wilcoxon rank test). Empty circles indicate results using all introns passing the initial filter criteria, while full circles show results for introns with consistent effects across replicates. **(b)** Survival curves of worms with *ama-1 (m322)* mutation (**left, replicate 1**) and flies with *RpII215*^*C4*^ mutation (**right, averaged survival curve**); worms 4 replicates, flies 3 replicates. Animals with slow Pol-II have a significantly increased lifespan (+20 % and +10 % median lifespan increase for *C. elegans* (n=120, P < 0.001, log rank test) and *D. melanogaster* (n=220, P < 0.001, log rank test), respectively).

Optimal elongation rates are required for fidelity of alternative splicing (*22, 23)*: for some exons, slow elongation favors weak splice sites that lead to exon inclusion, while these exons are skipped if elongation is faster (*5, 24, 25)*. Faster elongation rates can also promote intron retention leading to the degradation of transcripts via nonsense-mediated decay (NMD) *(26)* and possibly contributing to disease phenotypes *(27)*. Therefore, we next quantified changes in splicing. The first measure we used was splicing efficiency, which is the fraction of spliced reads from all reads aligning to a given splice site *(28)*. In most datasets, from total and nascent RNA-seq, we observed an increase of the spliced exon junctions relative to unspliced junctions during aging, and a decrease of the percent spliced junctions under lifespan-extending conditions (**Fig. 3a)**. Consistent with earlier work *(29)*, we observed more spliced transcripts under conditions of increased Pol-II speed, i.e. greater splicing efficiency. For co-transcriptional splicing to occur, Pol-II first needs to transcribe all parts relevant to the splicing reaction (i.e., 5’ donor, branch point, 3’ acceptor), which are located at the opposite ends of an intron (30,31). Our data suggest that accelerated transcription shortens the interval in which splicing choices are made, thus shortening the time between nascent RNA synthesis and intron removal.

**Fig. 3:**
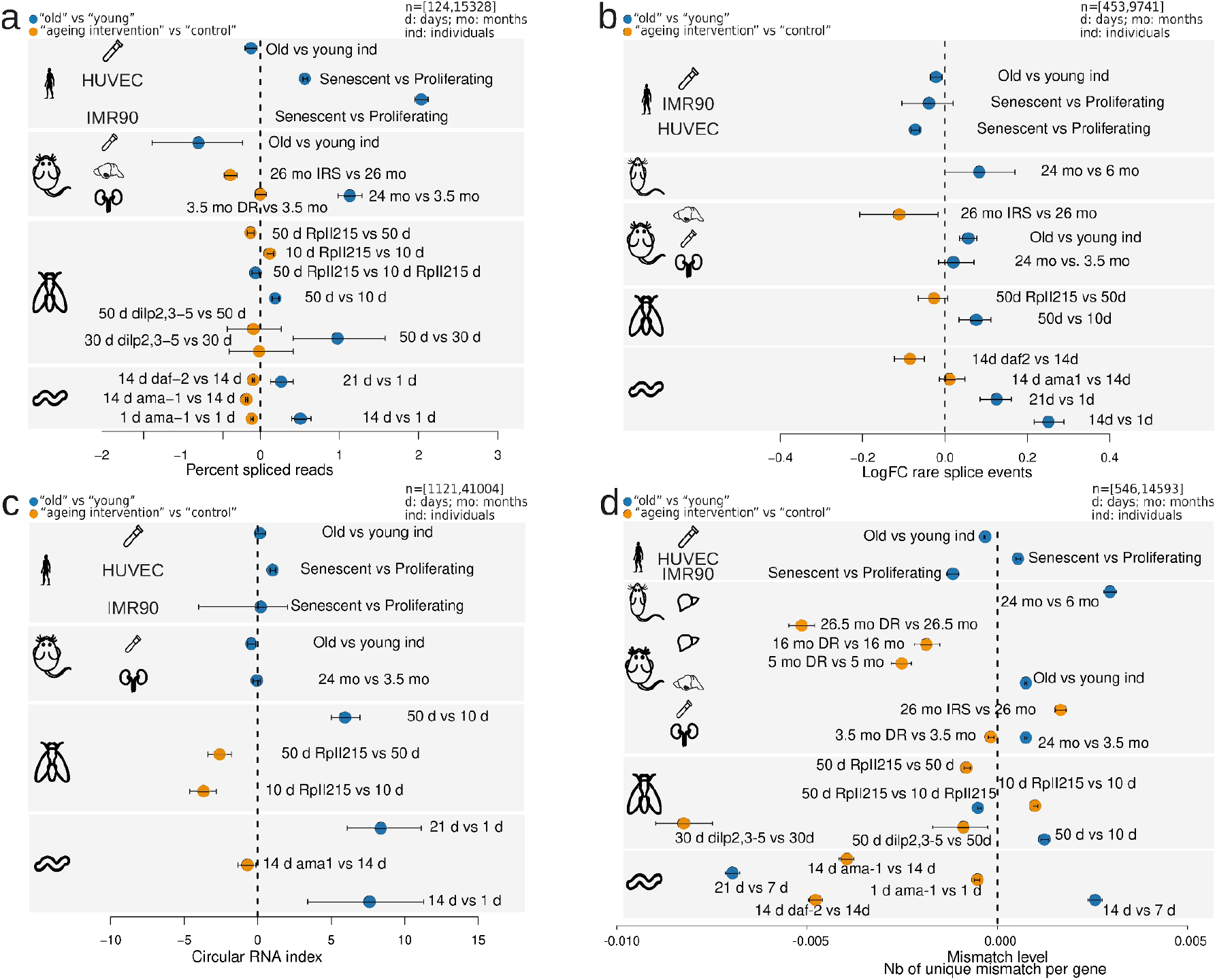
Changes in transcript structure upon ageing (old *vs*. young; blue) and after lifespan extending interventions (orange). Error bars show median variation ±95% confidence interval. **(a)** Average percent changes of rare splice events (<= 0.07 percent of total gene expression). Number of genes considered (*n*) ranged from 1121 to 41004. **(b)** Circular RNA index (back-spliced reads divided by sum of linear and back-spliced reads) for worms, fly heads, mouse and rat liver, human cell lines. Number of back-spliced junctions considered (*n*) ranged from 453 to 9741. **(c)** Average mismatch level changes. Number of genes considered (*n*) ranged from 546 to 14593. **(d)** Average changes of the fraction of spliced transcripts. Number of genes considered (*n*) ranged from 124 to 15328.

Accelerated transcription and splicing carries the risk of increasing the frequency of erroneous splicing events, which has been associated with advanced age and shortened lifespan *(32-35)*. It is non-trivial to deduce whether a specific splice isoform is the product of erroneous splicing or created in response to a specific signal. Simply checking if an observed isoform is annotated in some database can be problematic for multiple reasons. For instance, most databases have been created on the basis of data from young animals or embryonic tissue. Thus, a detected isoform that only may be functionally relevant in old animals will not be reported in such databases. Moreover, an annotated isoform might be the result of erroneous splicing if its expression is normally suppressed at a particular age or cellular context. We therefore based our analysis on the notion that extremely rare isoforms (rare with respect to all other isoforms of the same gene in the same sample) are more likely erroneous than frequent ones (*36,37)*. We used Leafcutter (*38*), which performs *de novo* quantification of exon-exon junctions based on split-mapped RNA-seq reads. Due to its ability to identify alternatively excised intron clusters Leafcutter is particularly suitable to study rare exon-exon junctions (*39)*. We defined rare splicing events as exon-exon junctions supported by ≤0.7% of the total number of reads in a given intron cluster, and the gene-specific fraction of rare clusters was computed as the number of rare exon-exon junctions divided by the total number of detected exon-exon junctions in that gene. We observed that such rare exon-exon junctions often resulted from exon skipping or from the usage of cryptic splice sites (**Extended Data Fig. 10**). The average fraction of rare splicing events increased during aging in fly and worm, and this effect was reverted under most lifespan-extending conditions (**Fig. 3b**). However, we did not observe a consistent age-associated increase of the fraction of rare splice variants across all species, which may at least in part be due to the more complex organization of splice regulation in mammalian cells.

Another potential indicator of transcriptional noise is the increased formation of circular RNAs (circRNAs) *(40,41), i*.*e*., of back-spliced transcripts with covalently linked 3’ and 5’ ends *(43)*. Increased Pol-II speed has previously been associated with increased circRNA abundance (*42*). Thus, we quantified the fraction of circRNAs as the number of back-spliced junctions normalized by the sum of back-spliced fragments and linearly spliced fragments. We observed either increased or unchanged average circRNA fractions during aging, while reducing Pol-II speed also reduced circRNA formation (**Fig. 3c**). This suggests that faster Pol-II elongation correlates with a general increase of circRNAs. Nevertheless, our data does not provide evidence that increased circRNA levels directly result from increased Pol-II speed, despite it being a consequence of the overall reduced quality in RNA production.

Increased Pol-II speeds can lead to more transcriptional errors, because the proofreading capacity of Pol-II is challenged (*8)*. To assess the potential impact of accelerated elongation on transcript quality beyond splicing, we measured the number of mismatches in aligned reads for each gene. For this, we normalized mismatch occurrence to individual gene expression levels and excluded mismatches that were likely due to genomic variation or other artifacts (see **Methods** for details). We observed that the average fraction of mismatches increased with age, but decreased under most lifespan-extending treatments (**Fig. 3d**). Consistent with prior findings (*8)*, slow Pol-II mutants exhibited reduced numbers of mismatches compared to wild-type control levels in 3 out 4 comparisons.

Subsequently, we explored alterations in chromatin structure as a possible cause of the age-associated changes in Pol-II speeds. Nucleosome positioning along DNA is known to affect both Pol-II elongation and splicing *(16,44-46)*. Furthermore, aged eukaryotic cells display reduced nucleosomal density in chromatin and ‘fuzzier’ core nucleosome positioning *(47,48)*. Thus, age-associated changes in chromatin structure could contribute to the changes in Pol-II speed and splicing efficiency that we observed. To test this, we performed micrococcal nuclease (MNase) digestion of chromatin from early (proliferating) and late-passage (senescent) human IMR90 cells, followed by ∼400 million paired-end read sequencing of mononucleosomal DNA (MNase-seq). Following mapping, we examined nucleosome occupancy. In senescent cells, introns were less densely populated with nucleosomes compared to proliferating cells *(49)* (**Fig. 4a**). In addition, we quantified peak ‘sharpness’, reflecting the precision of nucleosome positioning in a given MNase-seq dataset (see **Methods**), as well as the distances between consecutive nucleosomal summits as a measure of the spacing regularity (*49,50)*. Principle Component Analysis (PCA) of the resulting signatures indicated consistent changes of nucleosome ‘sharpness’ and distances upon entry into senescence as the samples clearly separated by condition **(Fig. 4b, c)**. Both measures were significantly, but moderately, altered in senescent cells (**Fig. 4d,e**): average sharpness was slightly decreased (along both exons and introns), and average inter-nucleosomal distances slightly increased in introns. In conclusion, the transition from a proliferating cell state to replicative senescence was associated with small, but significant changes in chromatin structure, involving nucleosome density and positioning-changes that were previously shown to affect Pol-II elongation *(44,48,51)*.

**Fig. 4:**
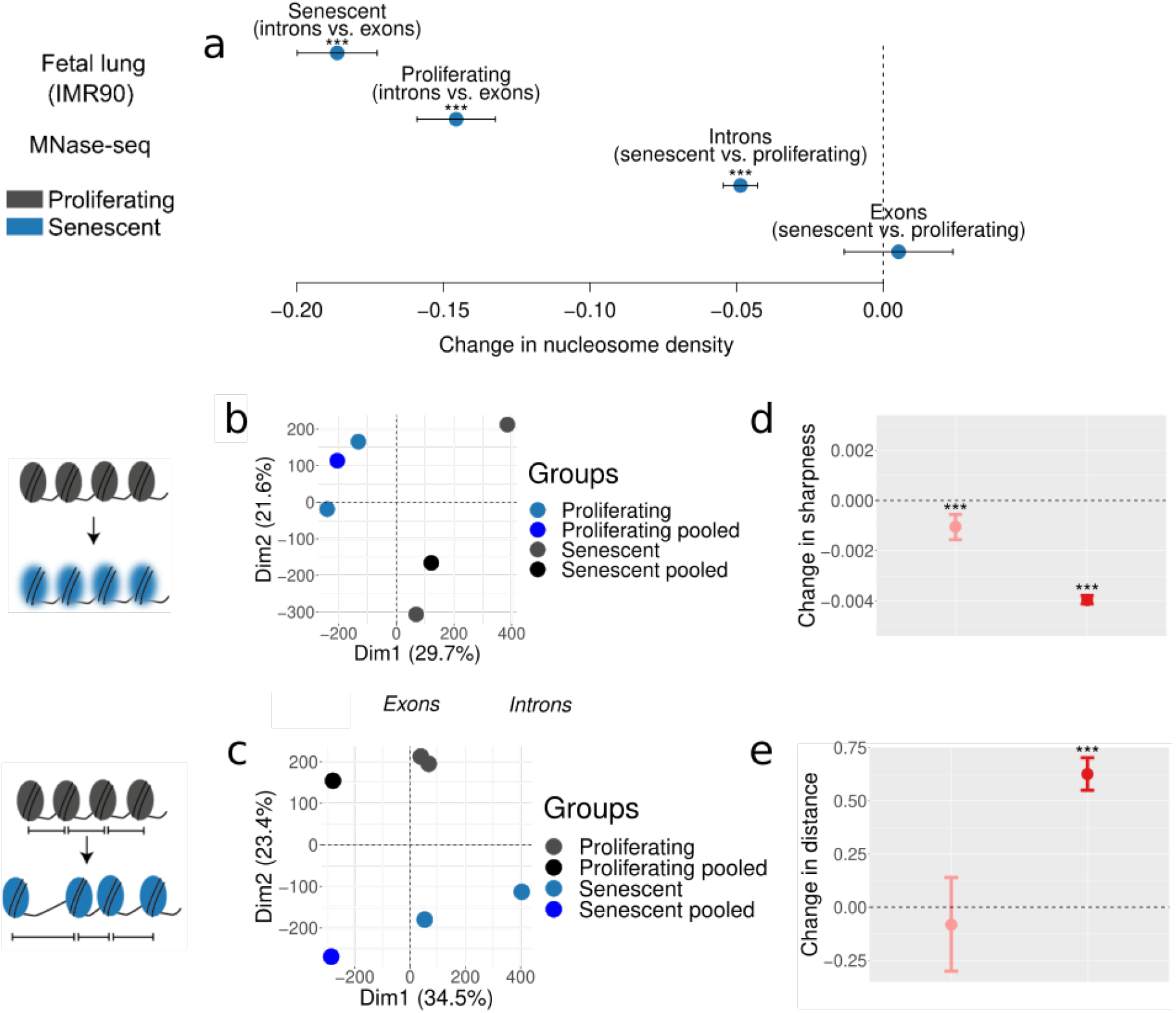
Profiling of nucleosome positioning in human cell models. **(a)** Average differences in nucleosome density between exons (n=37,625) and introns (n=193,912), and between proliferating and senescent cells. (**b**) Changes of nucleosome sharpness between senescent and proliferating cells in exons (left) and introns (right). (**d**) Distributions of distances between nucleosome summits between senescent and proliferating cells in exons (left) and introns (right). **(c+e)** PCA plots of nucleosome sharpness (**c)** and distances between nucleosome summits **(e)** in introns for individual samples and pooled data. All panels: Error bars show median variation ±95% confidence interval. Statistical significance of difference in pseudomedian distribution indicated by asterisks (*** P < 0.001, paired Wilcoxon rank test).

The organization of nucleosomes is severely influenced by histone availability *(46,47*). For example, histone H3 depletion reduces nucleosomal density and renders chromatin more accessible to MNase digestion (*52)*. Such global loss of histones constitutes a hallmark of aging and senescence *(53*). Consistent with this, our senescent IMR90 and HUVECs carry significantly reduced histone H3 protein levels (**Fig. 5a**). Conversely, elevated histone levels promote lifespan extension in yeast *(47*), *C. elegans (54)* and *D. melanogaster (56)*. To assess whether Pol-II elongation speed and senescence entry in human cells are causally affected by changes in nucleosomal density, we generated IMR90 populations homogeneously overexpressing GFP-tagged H3 or H4 in an inducible manner (**Fig. 5b** and **Extended Data Fig. 11a,b**). Overexpression of either histone resulted in significant reduction of Pol-II speed, confirming the causal connection between chromatin structure and transcriptional elongation (**Fig. 5c**). Pol-II speed reduction was accompanied by markedly reduced senescence-associated β-galactosidase staining in H3-/H4-overexpressing cells compared to both control (GFP-only) and uninduced cells (**Fig. 5d**). Moreover, both H3- and H4-overexpressing cells did not display p21 induction or HMGB1 depletion, both hallmarks of senescence entry, compared to control IMR90 (**Fig. 5e** and **Extended Data Fig. 11c**). Finally, MTT assays showed that viability and proliferation were improved in H3- and, to a lesser extent, in H4-overexpressing cells compared to control cells (**Fig. 5f**). Together, these results suggest that H3/H4 overexpression decelerates Pol-II and compensates for the aging-induced core histone loss *(45,51*) to restrict senescence entry.

**Fig. 5:**
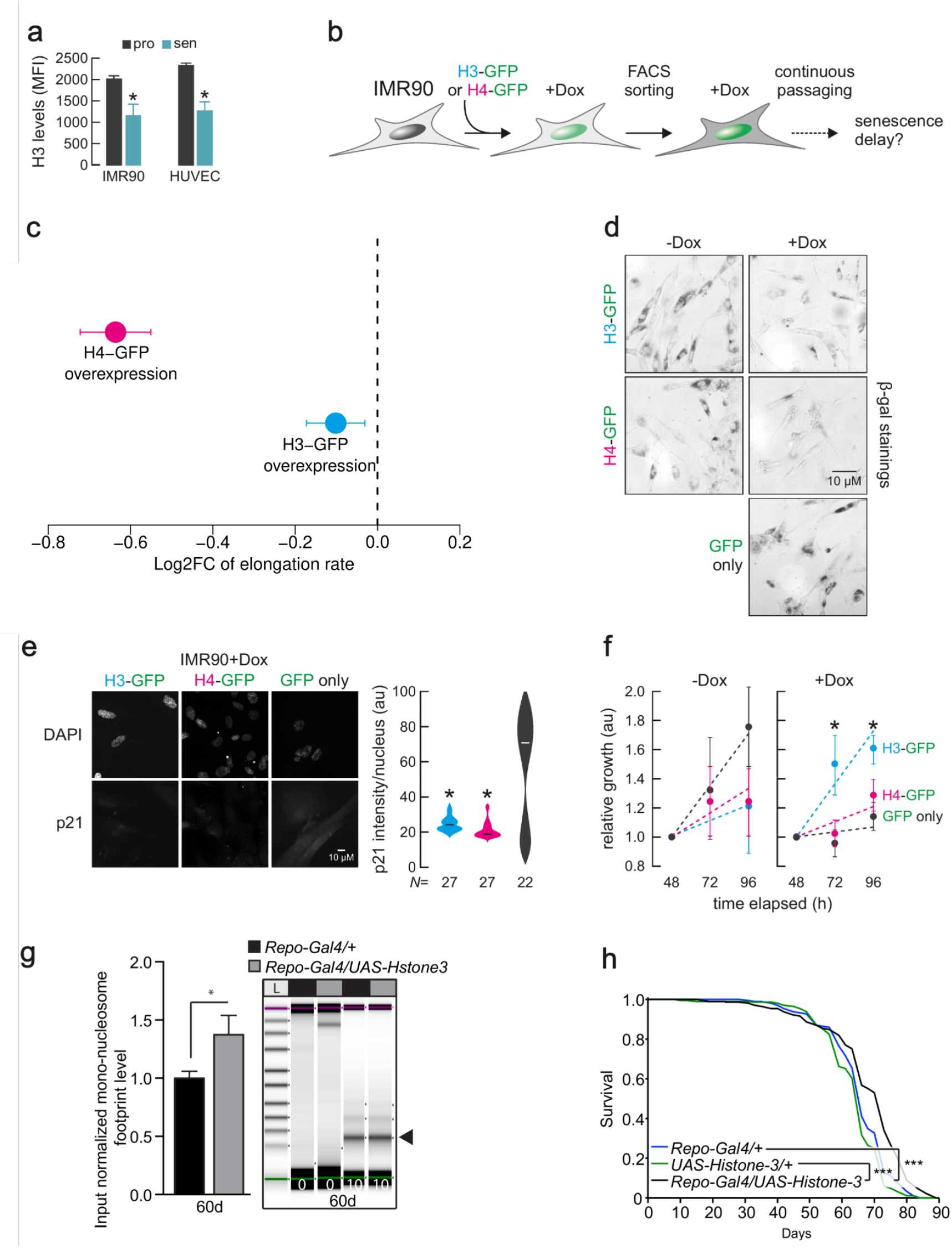
Histone overexpression slows down entry into senescence and decreases RNA-Pol-II speed. **(a)** H3 protein levels in proliferating and senescent HUVEC and IMR90 cells. **(b)** Schematic representation of the experiment. **(c)** Differences of average Pol-II elongation speeds between histone overexpression mutants and wild-type IMR90 cells (derived from 1,212 introns). Error bars show median variation ±95% confidence interval. All average changes of Pol-II elongation speeds are significantly different from zero (P < 0.01; paired Wilcoxon rank test). **(d)** Typical images from Beta-galactosidase(β-gal) staining of H3-GFP, H4-GFP and control IMR90 cells, in the presence and absence of Doxycycline. **(e)** Typical immunofluorescence images of H3-GFP, H4-GFP and control IMR90 cells (*left)* show reduced p21 levels in histone overexpression nuclei. Violin plots(*right*) quantify this reduction. N specifies the number of cells analyzed per condition. **(f)** MTT proliferation assay. **(g)** Quantification of input normalized mono-nucleosome footprints (black arrowhead) between the heads of aged (60d) flies overexpressing *Histone* 3 in glial cells (*Repo-Gal4/UAS-Histone 3*) and control fly heads (*Repo-Gal4/+*), significance determined by paired *t-*test (n>5, * *p<0*.*05*). Digests were halted after 10 min and visualized by Tapestation (Agilent) (n>5). **(h)** Lifespan analysis of flies *Repo-Gal4/UAS-Histone3* and control flies (*+/Repo-Gal4* and *UAS-Histone 3/+*) (n>100).

The average speed reduction following H4 overexpression was significantly larger than that obtained upon H3 overexpression, yet H4-overexpressing near-senescent IMR90 only marginally outperformed control cells in MTT assays (**Fig. 5f**). This raises the possibility of excessive reduction in Pol-II speed negatively affecting aspects of cell function *(55)*. To address the role of nucleosome density in organismal lifespan, we used UAS-His3 (*56*) to overexpress *His3*, specifically in *Drosophila* glial cells using Repo-Gal4 specifically in glial cells. H3 overexpression led to significantly increased numbers of mono-nucleosomes in aged (60 day-old) compared to the wild-type fly heads (**Fig. 5g**), thus possibly compensating for age-associated loss of histone proteins. Further, H3 overexpression in glial cells increased fruit fly lifespan (**Fig. 5h**). These *in vivo* results are consistent with our *in vitro* data from IMR90, demonstrating that H3 overexpression partially reverts the aging effects on chromatin density and promotes longevity in flies. As this was linked to a reversal in Pol-II elongation speed, our findings, together with earlier ones in yeast*(47,52), C. elegans (54)* and *D. melanogaster (56)*, demonstrate how the structure of the chromatin fiber likely modulates Pol-II elongation speed and lifespan.

## Discussion

We found a consistent increase in average intronic Pol-II elongation speed with age across four animal models, two human cell lines and human blood, and could revert this trend by employing lifespan-extending treatments. We also documented aging-related changes in splicing and transcript quality, such as elevated formation of circular RNAs and increased numbers of mismatches with genome sequences, which likely contribute to age-associated phenotypes. Further, we observed a consistent increase in the ratios of spliced to unspliced transcripts (splicing efficiency) with age across species (Fig. 3a), which has been reported to be a result of increased elongation speed *(29)*. However, we cannot exclude the possibility that this increase resulted from changes in RNA half-lives. Although average speed changes were predominantly significant, they remained small in absolute terms. This is expected, as drastic, genome-wide changes of RNA biosynthesis would quickly be detrimental for cellular functions and likely lead to early death. Instead, what we monitored here is a gradual reduction of cellular fitness characteristic for normal aging. Critically, we were able to increase lifespan in two species by decelerating Pol-II. Thus, despite being small in magnitude and stochastically emerging in tissues or cell populations, these effects are clearly relevant for organismal lifespan.

Genes exhibiting accelerated Pol-II elongation were not enriched for specific cellular processes, indicating that speed increase is probably not a deterministically cell-regulated response, but rather a spontaneous age-associated defect. Yet, the genes affected were not completely random, since we observed consistent changes across replicates for a subset of introns. Thus, there must be location-specific factors influencing which genomic regions are more prone to Pol-II speed increase and which not. This observation is consistent with earlier findings and our data, indicating that chromatin structure may causally contribute to age-associated Pol-II speed increase. Although we still lack a complete understanding of the molecular events driving Pol-II speed increase, our findings indicate that aging-associated changes in chromatin structure play an important role.

Our work establishes Pol-II elongation speed as an important contributor to molecular and physiological traits with implications beyond aging. Misregulation of transcriptional elongation reduces cellular and organismal fitness and may therefore contribute to disease phenotypes *(54,57,58,59)*. Taken together, the data presented here reveal a new molecular mechanism contributing to aging and serve as a means for assessing the fidelity of the cellular machinery during aging and disease.

## Methods

### Worm strains and demography assays

Nematodes were cultured using standard techniques at 20 **°** C on NGM agar plates and were fed with *E*.*coli* strain OP50. *DR786* strain carrying the *ama-1(m322)* IV mutation in the large subunit of Pol-II (RBP1), which confers alpha-amanitin resistance, was obtained from Caenorhabditis Genetics Center (CGC) *(59, 60)*. DR786 strain was then outcrossed into *N2 wild type* strain 4 times and mutation was confirmed by sequencing. 5’-3’ AGAAGGTCACACAATCGGAATC primer was used for sequencing. For each genotype, minimum of 120 age-matched day 1 young adults were scored every other day for survival and transferred to new plates to avoid starvation and carry-over progeny. Lifespan analyses using the C. elegans Lifespan Machine were conducted as previously described *(61)*. Briefly, wild-type N2 and mutant worms were synchronized by egg-prep (hypochlorite treatment) and grown on NGM-agar plates seeded with OP50 at 20°C. Upon reaching L4 stage these worms were transferred onto plates containing 0.1 g/ml 5-Fluoro-2′-deoxyuridine (FUDR) and placed into the modified flatbad scanners (35 Worms per plate). The scan interval was 30 min. Objects falsely identified as worms were censored. Time of death was automatically determined by the C. elegans Lifespan Machine *(61)*. Demography experiments were repeated multiple times. For all experiments, genotypes were blinded. Statistical analyses were performed using Mantel-Cox log rank method.

### Measurements of pharyngeal pumping rates in worms

Synchronized wild type and ama-1 (m322) animals were placed on regular NGM plates seeded with OP50 bacteria on day1 and day8 adulthood and number of pharyngeal pumping rate was assessed by observing the number of pharyngeal contractions during a 10sec interval using dissecting microscope and Leica Application Suite X imaging software. Pharyngeal pumping rate was then adjusted for number of pharyngeal pumping per minute. Animals that displayed bursting, internal hatching and death were excluded from the experiments. Experiments were repeated three independent times in a blinded fashion, scoring minimum of 15 randomly selected animals per genotype and time point for each experiment. One-way Anova with Tukey’s multiple comparison test was used for statistical significance testing. p-value < 0.0001****, error bars represent standard deviation.

### Fly strains and fly maintenance

The *RpII215*^*C4*^ fly strain (RRID:BDSC_3663), which carries a single point mutation (R741H) in the gene encoding the *Drosophila* RNA polymerase II 215kD subunit (RBP1), was received from the Bloomington *Drosophila* Stock Center (Bloomington, Indiana, USA). Flies carrying the *RpII215*^*C4*^ allele *(62)* are homozygous viable but show a reduced transcription elongation rate *(19). RpII215*^*C4*^ mutants were backcrossed for 6 generations into the outbred *white Dahomey (wDah)* wild type strain. A PCR screening strategy was used to follow the *RpII215*^*C4*^ allele during backcrossing. Therefore, genomic DNA from individual flies was used as a template for a PCR reaction with primers SOL1064 (CCGGATCACTGCTGCATATTTGTT) and SOL1047 (CCGCGCGACTCAGGACCAT). The 582 *bp* PCR product was restricted with BspHI, which specifically cuts only in the *RpII215*^*C4*^ allele, resulting in two bands of 281 *bp* and 300 *bp*. At least 20 individual positive female flies were used for each backcrossing round. Long-lived insulin mutant flies, which lack three of the seven *Drosophila* insulin-like peptides, *dilp2-3,5* mutants (RRID:BDSC_30889) *(63*, were also backcrossed into the *wDah* strain, which was used as wild type control in all fly experiments. Flies were maintained and experiments were conducted on 1,0 SY-A medium at 25°C and 65% humidity on a 12L:12D cycle *(63)*.

### Fly lifespan assays

For lifespan assays, fly eggs of homozygous parental flies were collected during a 12 h time window and the same volume of embryos was transferred to each rearing bottle, ensuring standard larval density. Flies that eclosed during a 12 h time window were transferred to fresh bottles and were allowed to mate for 48h. Subsequently, flies were sorted under brief CO_2_ anesthesia and transferred to vials. Flies were maintained at a density of 15 flies per vial and were transferred to fresh vials every two to three days and the number of dead flies was counted. Lifespan data were recorded using Excel and were subjected to survival analysis (log rank test) and presented as survival curves.

### Mouse maintenance and dietary restriction protocol

The DR study was performed in accordance with the recommendations and guideline of the Federation of the European Laboratory Animal Science Association (FELASA), with all protocols approved by the Landesamt für Natur, Umwelt und Verbraucherschutz, Nordrhein-Westfalen, Germany. Details of the mouse liver DR protocol have been published previously (Hahn et al., 2017). For the mouse kidney, male *C57BL/6* mice were housed under identical SPF conditions in group cages (5 or fewer animals per cage) at a relative humidity of 50-60% and a 12 hour light and 12 hour dark rhythm. For dietary restriction vs control, 8 week old mice were used. Dietary restriction was applied for 4 weeks. Control mice received food and water ad libitum. Mice were sacrificed at 12 weeks. For comparison of young vs aged mice, 14 week and 96 week old mice were used. Food was obtained from ssniff (Art. V1534-703, Soest, Germany) and Special Diet Services (Witham, UK). The average amount of food consumed by a mouse was determined by daily weighing for a period of two weeks and was on average 4,3 g per day. DR was applied for 4 weeks by feeding 70% of the measured ad libitum amount of food. Water was provided ad libitum. Mice were weighed weekly to monitor weight loss. Neither increased mortality nor morbidity was observed during dietary restriction.

### RNA extraction

Wild-type N2 strain, alpha-amanitin resistant *ama-1(m322)* mutants and long-lived insR/IGF signaling mutants, *daf-2*(*e1370*) were sent for RNA-seq. For each genotype, more than 300 aged-matched adult worms at desired time points were collected in Trizol (Thermo Fisher Scientific, USA) in 3 biological replicates. Total RNA was extracted using RNAeasy Mini kit (Qiagen, Hilden, Germany). The RNA-seq data for brains of 30 days and 50 days old *dilp2-3,5* and *wDah* control flies have been published previously (Weigelt et al., 2020, Molecular Cell). 10 days and 50 days old *RpII215*^*C4*^ mutants and *wDah* control flies were snap frozen and fly heads were isolated by vortexing and sieving on dry ice. Total RNA from three biological samples per treatment group was prepared using Trizol Reagent according to the manufacturer’s instructions, followed by DNAse treatment with the TURBO DNA-free Kit (Thermo Fisher Scientific). Mouse liver samples were isolated from 5, 16 and 27 months old ad libitum and DR animals, which corresponded to 2, 13 and 24 months of DR treatment, respectively. RNA was isolated by Trizol and DNase-treated. The RNA-seq data for 5 and 27 months old liver DR samples have been published previously *(64)*, while the 16 months data are first published here. RNeasy mini Kit and Trizol were used to isolate RNA from snap-frozen kidneys as per manufacturer’s instructions. Hypothalamus tissue of long-lived insulin receptor substrate 1 (*IRS1-/-*) knock out mice (65) and *C57BL/6* black control animals was dissected manually at the age of 27 months. RNA was isolated by Trizol with subsequent DNase treatment. For blood samples globin RNA was removed using GLOBINclear™ Kit, mouse/rat/human, for globin mRNA depletion.

### Human whole blood sample acquisition and RNA extraction

Human samples were obtained as part of a clinical study on aging-associated molecular changes (German Clinical Trials Register: DRKS00014637) at University Hospital Cologne. The study cohort consisted of healthy subjects between 21 and 70 years of age. Whole blood samples were obtained using the PAXgene Blood RNA system (Becton Dickinson GmbH, Heidelberg, Germany) directly after informed consent. After storage at -80°C for at least 24 h RNA extraction was performed by usage of PAXgene Blood RNA Kit (Quiagen, Hilden, Germany) according to the manufacturer’s protocol. The study was operated in accordance with the Declaration of Helsinki and the good clinical practice guidelines by the International Conference on Harmonization. All patients provided informed consent and approval of each study protocol was obtained from the local institutional review board (Ethics committee of the University of Cologne, Cologne, Germany; (17-362, 2018-01-17).)

### Human cell culture

Human umbilical vein endothelial cells (HUVECs) from pooled donors (Lonza, Cologne, Germany) and Human fetal lung (IMR90) cells (from two donors) were grown to 80% to 90% confluence in endothelial basal medium 2-MV with supplements (EBM; Lonza) and 5% fetal bovine serum (FBS) and MEM (Sigma-Aldrich) with 20 FBS (Gibco) and 1% non-essential amino acids (Sigma-Aldrich) for HUVECs and IMR-90 respectively.

### Total RNA and nascent RNA sequencing

From 1 µg input of total RNA, ribosomal RNA was removed using the Ribo-Zero Human/Mouse/Rat kit (Illumina). Sequencing libraries were generated according to the TruSeq stranded total RNA (Illumina) protocol. To generate the final cDNA library, products were purified and amplified by PCR for 15 cycles. After validation and quantification of the library on an Agilent 2100 Bioanalyzer, equimolar amounts of libraries were pooled. Pools of 5 libraries were sequenced per lane on an Illumina HiSeq 4000 sequencer. For a description of all the RNA-seq datasets used in this study see Extended Data Table 1. The same protocol was used to sequence cDNA libraries from human cell “factory” RNA, which was isolated as described previously (66).

### RNA-seq alignments and gene expression analysis

Raw reads were trimmed with trimmomatic version 0.33 (67) using parameters ‘ILLUMINACLIP:./Trimmomatic-0.33/adapters/TruSeq3-PE.fa:2:30:10 LEADING:3 TRAILING:3 SLIDINGWINDOW:4:15 MINLEN:45’ for paired-end datasets and ‘ILLUMINACLIP:./Trimmomatic-0.33/adapters/TruSeq3-SE.fa:2:30:10 LEADING:3 TRAILING:3 SLIDINGWINDOW:4:15 MINLEN:45’ for single-end datasets. Alignment was performed with STAR version 2.5.1b (68) using the following parameters: ‘--outFilterType BySJout --outWigNorm None’ on the genome version mm10, rn5, hg38, dm6, ce5 for *M. musculus, R. norvegicus, H. sapiens, D. melanogaster, C. elegans*, respectively. We estimated transcript counts using Kallisto version 0.42.5 for each sample. To determine differentially expressed genes we used DESeq2 version 1.8.2 (69) with RUVr normalization version 1.6.2 (70). For the differential analysis of transcriptional elongation regulators, we downloaded the list of positive and negative regulators from the GSEA/MSigDB (71). Gene ontology (GO) term enrichment analysis of differentially expressed genes or genes with increased RNA-Pol-II elongation speed was carried out using TopGO version 2.20.0. For GO enrichment analysis of differentially expressed genes, we identified 4784 genes as evolutionarily conserved from each species of our study to humans: genes were either direct orthologues (one2one) or fusion genes (one2many) of *H. sapiens* were retrieved from ENSEMBL database using biomaRt 2.24.1 (72). Using our 4784 genes evolutionary conserved, we further divided into consistently up-regulated or down-regulated genes across species during aging or ‘aging intervention’ (as target set for GO enrichment: aging up-regulated: 92 genes; aging down-regulated: 71; ‘aging intervention’ up-regulated: 164 genes; ‘aging intervention’ down-regulated: 473 genes; as background set 4784 orthologue genes between *R. norvegicus, M. musculus, D. melanogaster, C. elegans*, and *H. hsapiens*). For GO enrichment analysis of genes harboring increasing Pol-II speed, we used as target set the top 200 or 300 genes with an increase in Pol-II speed change for each species. Quantification of transcript abundance for ITPR1 and AGO3 was obtained by using StringTie(73). For circular RNA, we aligned the reads using STAR version 2.5.1b (68) with the following parameters:’ --chimSegmentMin 15 --outSJfilterOverhangMin 15 15 15 15 --alignSJoverhangMin 15 --alignSJDBoverhangMin 15 --seedSearchStartLmax 30 -- outFilterMultimapNmax 20 --outFilterScoreMin 1 --outFilterMatchNmin 1 -- outFilterMismatchNmax 2 --chimScoreMin 15 --chimScoreSeparation 10 -- chimJunctionOverhangMin 15’. We then extracted back spliced reads from the STAR chimeric output file and normalized the number of back spliced reads by the sum of back spliced (*BS*_*i*_) and spliced reads from linear transcripts (*S1*_*i*_, *S2*_*i*_) for an exon *i (8)*:

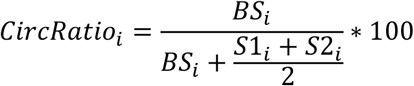

Here, *S1*_*i*_ refers to the number of linearly spliced reads at the 5’ end of the exon and *S2*_*i*_ refers to the respective number of reads at the 3’ end of the exon. Thus, this score quantifies the percent of transcripts from this locus that resulted in circular RNA. Finally, we quantified the significance of the average change in circular RNA formation between two conditions using the Wilcoxon rank test.

### Definition of intronic regions

All annotation files for this analysis were downloaded from the Ensembl website (74) using genome version ce10 for *Caenorhabditis elegans*, mm10 for *Mus musculus*, hg38 for *Homo sapiens*, rn5 for *Rattus norvegicus*, and dm6 for *Drosophila melanogaster*. The following filtering steps were applied on the intronic ENSEMBL annotation files: First we removed overlapping regions between introns and exons in order to avoid confounding signals due to variation in splicing or transcription initiation and termination. Overlapping introns were merged to remove duplicated regions from the analysis. In the next step we used STAR (68) to detect splice junctions and compared them to the intronic regions. Introns with at least 5 split reads bridging the intron (i.e. mapping to the flanking exons) per condition were kept for subsequent analyses. Thereby we ensured a minimum expression level of the spliced transcript. When splice junctions were detected within introns, we further subdivided those introns accordingly. Introns with splice junction straddling were discarded. The above-mentioned steps were performed using Bedtools version 2.22.1 using subtract and merge commands. After these filtering steps, the number of usable introns per sample varied between a few hundred (*n*=546, *C. elegans*, total RNA) to over ten thousand (*n*=13,790, *H. sapiens*, nascent-RNA-seq). These large differences resulted from different sequencing depths, sequencing quality (number of usable reads), and from the complexity of the genome (numbers and sizes of introns, number of alternative isoforms, etc.). In order to avoid artifacts due to the different numbers of introns used per sample, we always contrasted the same sets of introns for each comparison of different conditions (e.g. old *versus* young, treatment *versus* control). Note that certain comparisons were not possible for all species, due to variations in the experimental design. For instance, for mouse kidney only a single time point after lifespan intervention (dietary restriction DR, age 3 months) was available, which prevented a comparison of old versus young DR mice, but allowed comparison with *ad libitum* fed mice at the young age.

### Transcriptional elongation speed based on intronic read distribution

In order to calculate Pol-II speeds we used RNA-seq data obtained from total RNA *(75)* and nascent RNA (65, 76) enrichment. In contrast to the widely used polyA enrichment method *(77)*, which primarily captures mature, spliced mRNAs and is therefore not suitable to estimate Pol-II speeds based on intronic reads, these methods yield sufficient intronic coverage to quantify elongation rates. To analyze the distribution of intronic reads between conditions, we devised a score for each intron. We fitted the read gradient (slope) along each of selected introns (5’->3’; see above for the filtering criteria). Note that the intron gradient is not influenced by exonucleolytic degradation of excised intron lariats *(78)* and that this measure is only weakly associated with the expression level of the transcript *(17)* (Extended Data Table 1).

In order to transform slopes to Pol-II elongation speed we used the following formalism. We assume an intron of length *L* and we assume that at steady state a constant number of polymerases is initiating and the same number of polymerases is terminating at the end of the intron; i.e. we assume that premature termination inside the intron can be ignored. Polymerases are progressing at a common speed of *k* [*bp/min*]. The average time that it takes a polymerase to traverse the whole intron is hence

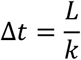

Transcription is initiated at a rate of *n* polymerases per unit time [*1/min*]. Hence, the number of polymerases *N* initiating during Δ*t* is:

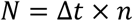

The slope s is the number of transcripts after the distance *L* minus the number of transcripts at the beginning divided by the length of the intron:

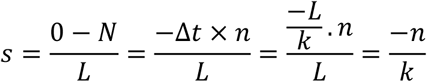

and thus, the speed *k* can be computed from the slope as:

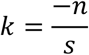

Hence, slope and speed are inversely related and the speed depends also on the initiation rate (i.e. the expression rate). However, we observed empirically only a small dependency between expression and slope *(17)* (Extended Data Table 1).

To validate our estimates of Pol-II speeds we compared our data with experimental values estimated via GRO-seq *(16)* and tiling microarray data *(12)*. There was a significant correlation (GRO-seq: R=0.38, p-value=4e-5, compared to time point 25-50 min see Jonkers et al. *(16)*; Tilling array; R=0.99, p-value=<2.6e-16, data not shown) between our data and experimentally measured transcriptional elongation values. We noted that our Pol-II speed estimates for different introns of the same gene were more similar than Pol-II speed estimates for random pairs of introns, implying that gene-specific factors or local chromatin structure influence Pol-II speed.

### 4sU-DRB labelling, TUC-conversion and elongation rate calculation

The estimation of transcription elongation speed using RNA labelling was based on the measurement of nucleotides added per time unit in a newly synthesized nascent transcript. First, transcription was reversibly inhibited by 6-dichlorobenzimidazole 1-β-d-ribofuranoside (DRB) in order to achieve accumulation of RNA polymerase II at the transcription start sites and synchronized transcriptional elongation initiation upon DRB removal. Simultaneously with the DRB removal, cells were pulsed for different time points with the Uridine-analogue 4-thiouridine (4sU) in order to enrich for newly synthesized transcripts. Finally, total RNA was isolated per each time point and the RNA polymerase II speed was determined by calculating the 4sU nucleotides added to the nascent transcript per time point. To estimate RNA polymerase II speed change in aging cells, human fetal lung fibroblasts (IMR90) in proliferating and in senescent state were treated using this experimental procedure.

In order to select the time-points to be used in the experiment, validate the DRB treatment and removal and check the enrichment efficiency of 4sU, a control experiment was set according to the protocol of Fuchs *et al*. 2015 (79). Two million proliferating cells (passage 14) were treated with 100 µM DRB (Merck, D1916) in their medium for three hours at 37 °C and, upon DRB removal, they were pulsed with 1 mM 4sU (Sigma-Aldrich, T4509) for 0 min, 5 min, 15 min, 30 min, 45 min, 60 min, 90 min and 120 min. Immediately after the completion of each time point, cells were lysed in TRIzol (ThermoFisher, 15596018) and RNA was isolated with the Direct-Zol RNA mini-prep kit (ZymoResearch, R2052). To validate DRB treatment, qRT-PCR was performed in cDNA from all time points using the primers designed by Fuchs *et al*. 2015 (79) in proximal and distant introns of the OPA-1 gene. Furthermore, to estimate 4sU enrichment, the RNA collected in each time point was biotinylated using the EZ-Link biotin HPDP kit (ThermoFisher, 21341) and biotinylated RNA was enriched with streptavidin-coated beads (DYNAL™ Dynabeads™ M-280 Streptavidin, ThermoFisher 11205D). qRT-PCR evaluation was performed also with the primers suggested by Fuchs *et al*. 2015 *(79)* against TTC-17 nascent and mature mRNA and 18S rRNA.

For the actual experiment, we performed the Thiouridine-to-Cytidine Conversion Sequencing (TUC-Seq) protocol developed by Lusser *et al*. (2020) *(80)* in order to detect the 4sU labelled transcripts in different time points. In this method, the thiol group of 4sU is quantitatively converted to cytidine via oxidation by OsO_4_ in aqueous NH_4_Cl solution. The OsO_4_-treated RNA samples are submitted to RNA-sequencing to quantify labelled and non-labelled transcripts and define the number of reads containing Uridine-to-Cytidine conversions. To this aspect, nine million proliferating (passage 9) cells and nine million cells that had entered senescence (passage 35) were treated with 100 µM DRB for three hours at 37 °C. Immediately after DRB removal, cells were pulsed with 1 mM 4sU for 0 min, 5 min, 15 min, 30 min and 45 min. RNA was isolated manually according to the TRIzol protocol and treated with 40 Units DNAse I (ZymoResearch, E1010) for 20 min at room temperature. RNA was purified with the RNA Clean & Concentrator-25 kit (ZymoResearch, R1018) and quantified using a NanoDrop spectrophotometer. For the TUC-conversion, 10 µg of 4sU labelled RNA was treated with 1.43 mM OsO_4_ (Merck, 251755) in 180 mM NH_4_Cl (Merck, 09718) solution pH 8.88 for 3 hours at 40 °C as described in Lusser *et al*. (2020) *(80)*. Subsequent sample concentration and purification were also done according to this protocol. 4sU-labelled and OsO_4_-treated RNA samples derived from proliferating and senescent IMR90s in all five time-points were subjected to RNA-sequencing. As a negative control for the TUC-conversion, we used a mixture 1:1 of 4sU-labelled but not OsO_4_-treated samples from the time-points 30 min and 45 min. The RNA-sequencing was performed in two biological replicates per condition.

Detection of labelled transcripts was performed based on the Lusser protocol, modified for Illumina RNA-SEQ:

1. FASTQ files were aligned to the genome using STAR to produce BAM files.
2. Sam2tsv *(81)* was then used to identify single nucleotide mismatches.
3. A custom R script was used to count the number of A-G or T-C mismatches per read.

Only read-pairs with at least 3 A-G or T-C mismatches were assumed to be 4sU-labeled and thus retained for subsequent analyses. Because the 5 min samples contained a very low number of reads with conversions, they were discarded from the rest of the analysis. We employed two approaches for estimating the elongation rate per gene from the 4sU-labelling data. For the first approach, we tracked the progress of RNA-Pol-II complexes constructing single gene coverage profiles using 4sU-labeled reads. Progression was determined by picking the 99^th^ percentile of gene body coverage in each sample to determine the front of elongating RNA polymerases. (We did not use the last converted read to determine the front, because this measure would be too sensitive to noise in the data.) Elongation rates were calculated by fitting a linear model on the front positions of Pol II in 0, 15, 30 and 45 min in the first 100 kb of each gene. To determine elongation rates with greater accuracy, we filtered out genes with a length of less than 100 kb, since short genes can be fully transcribed in less than 45 min or even 30 min. This first approach of estimating RNA-Pol-II speeds is characterized by high accuracy, but is limited to genes longer than 100 kb. The data in Panel e of Figure 1 is based on this approach.

The direct comparison of the 4sU data to the approach using read-coverage slopes in introns required a large set of genes for which RNA-Pol-II speed could be measured using both assays. In order to maximize this gene set we devised a second alternative approach for deriving speed from 4sU-labelling data that is applicable to shorter genes. For this second approach we measured the front position of the polymerase in the same way as before (using the 99^th^ percentile) but across the whole gene. For genes 30kb to 100kb long, we calculated the elongation rate from the difference in the front positions of the polymerase at 15 mins and 30 mins and divided this distance by 15 minutes in order to obtain speed measures per minute. For genes more than 100kb long, we calculate the elongation rate from the difference in the positions of the polymerase at 30 mins and 45 mins divided by 15 minutes. This second speed measure is less accurate than the first one, because it uses only two time points per gene; however, it enables estimating speed for genes shorter than 100 kb. The data in Panel d of Figure 1 is based on this second measure. Note that both measures confirmed the increase in average RNA-Pol-II elongation speed from proliferating to senescent IMR90 cells.

### ^35^S-methionine/^35^S-cysteine incorporation to measure translation rates in Drosophila

E*x-vivo* incorporation of radio-labeled amino acids in fly heads was performed as previously described *(82)*. Briefly, 25 heads of each young (10 days) and old (50 days) wDah control and RpII215^C4^ mutant animals were dissected in replicates of 5 and collected in DMEM (#41965-047, Gibco) without supplements, at room temperature. For labeling, DMEM was replaced with methionine and cysteine free DMEM (#21-013-24, Gibco), supplemented with ^35^S-labeled methionine and cysteine (#NEG772, Perkin-Elmer). Samples were incubated for 60 min at room temperature on a shaking platform, then washed with ice cold PBS and lysed in RIPA buffer (150 mM sodium chloride, 1.0% NP-40, 0.5% sodium deoxycholate, 0.1% SDS, 50 mM Tris, pH 8.0) using a pestle gun (VWR, Germany). Lysates were centrifuged at 13.000 rpm and 4 °C for 10 min and protein was precipitated by adding 1 volume of 20% TCA, incubating for 15 min on ice and centrifugation at 13.000 rpm, 4 °C for 15 min. The pellet was washed twice in acetone and resuspended in 200 µl of 4 M guanine-HCl. 100µl of the sample was added to 10ml scintillation fluid (Ultima Gold, Perkin-Elmer) and counted for 5 min per sample in a scintillation counter (Perkin-Elmer). Protein determination was done in duplicates (25µl each) per sample using the Pierce BCA assay kit (Thermo Fisher Scientific). Scintillation counts were normalized to total protein content.

### Mismatch detection

Mismatch detection was performed using the tool rnaseqmut (https://github.com/davidliwei/rnaseqmut), which detects mutations from the NM tag of BAM files. To avoid detection of RNA editing or DNA damage-based events we only considered genomic positions with only 1 mismatch detected (i.e. occurring in only one single read). Reads with indels were excluded and only mismatches with a distance of more than 4 from the beginning and the end of the read were considered. A coverage level filter was applied so that only bases covered by at least 100 reads were kept. A substantial number of mismatches may result from technical sequencing errors. However, since young and old samples were always handled together in the same batch, we can exclude that consistent differences in the number of mismatches are due to technical biases. The fraction of RNA editing events is generally relatively low and not expected to globally increase with age *(83)*.

### MNase-seq sample preparation

Mononucleosomal DNA from proliferating and senescent IMR-90 cells (from two donors) were prepared and sequenced on an Illumina HiSeq4000 platform as previously described *(84)*. For fly heads, a MNase digestion assay was performed using the EZ nucleosomal DNA prep kit, as per the manufacturer’s guidelines (Zymo Research). Briefly, 25 snap frozen heads were lysed in nuclei prep buffer, and incubated on ice for 5min. Cuticle fragments were then removed via centrifugation (30sec 50xg). Nuclei were pelleted (5min 500xg) and washed twice in digestion buffer and resuspended in 100μl of digestion buffer. Nucleosome footprints were then digested using 0.05U of MNAse (Zymo Research). Samples were taken at 0, 2, 3 and 5min or 10 min for prolonged digestion, and immediately stopped in MN stop buffer (Zymo Research). Samples were isolated using Zymo Spin IIC columns. Nucleosome footprints (1:10 dilution) were visualized by Tapestation using High sensitivity D1000 ScreenTape (Agilent).

### MNase-seq analysis

We used nucleR *(85)* with default parameters to calculate peak sharpness as a combination of peak width and peak height. Peak ‘width’ was quantified as the standard deviation around the peak center and peak ‘height’ was quantified as the number of reads covering each peak *(85)* and the distance between peak summits. Intron and exon annotations were downloaded from UCSC table utilities *(74)* and filtered as described in Definition of intronic regions. Nucleosome density (Figure 5a) is defined as the number of nucleosome peaks found within an exon or an intron divided by the length of the exon or intron.

### Western blotting

Western blots were carried out on protein extracts of individual dissected tissues. Proteins were quantified using BCA (Pierce). Equal amounts were loaded on Any-KD pre-stained SDS-PAGE gels (Bio-Rad) and blotted according to standard protocols. Antibody dilutions varied depending on the antibody and are listed here: Histone 3 (1:1000), HP1 (DSHB) (1:500). Appropriate secondary antibodies conjugated to horseradish peroxidase were used at a dilution of 1:10000.

### Inducible histones overexpression

Doxycycline (Dox)-inducible expression of histones H3 and H4 in proliferating human fetal lung fibroblasts (IMR90) was achieved using the PiggyBac transposition system *(86)*. The open reading frames of H3 and H4 were cloned in the Dox-inducible expression vector KA0717 (KA0717_pPB-hCMV*1-cHA-IRESVenus was a gift from Hans Schöler, Addgene plasmid #124168; http://n2t.net/addgene:124168; RRID:Addgene_124168) fused at their 3’ end in frame to the sequence of the yellow fluorescent protein (YFP) mVenus *(87)*. After sequencing validation, each construct was co-transfected in IMR90s with the transactivator plasmid KA0637 (KA0637_pPBCAG-rtTAM2-IN was a gift from Hans Schöler, Addgene plasmid #124166; http://n2t.net/addgene:124166; RRID:Addgene_124166) and the Super piggyBac Transposase expression vector (SBI System Biosciences, PB200PA-1,) using the Lipofectamine™ LTX Reagent with PLUS™ Reagent (ThermoFisher Scientific, 15338100,) according to the manufacturer’s instruction. In total 2.5 µg of the vectors KA0717, KA0637 and PB200PA-1 were used for each transfection in 10:3:1 ratio. Stable transgene-positive cells were selected using 250 μg/ml G418 (resistance gene carried in KA0637) for 7 days. Emerging cells were induced for 24 h with 2.5 µg/ml doxycycline and then subjected to Fluorescent-Activated Cell Sorting (FACS) to select the ones expressing the mVenus (BD FACSAria™ II, BD Biosciences). H3 and H4 overexpression was verified by Western Blot with anti-H3 and anti-H4 antibodies (Abcam, ab1791 and ab10158 respectively). All further assays were repeated in proliferating cells and cells at the senescence entry state. Senescence state was monitored by Beta-galactosidase staining (88) in different passages (Cell Signaling Technology, Senescence β-Galactosidase Staining Kit, 9860). Immunofluorescence stainings (IF) to detect HMGB1, p21 and HMGB2 (Abcam, ab18256, ab184640 and ab67282 respectively) were performed as previously described (89) and images were acquired in a widefield Leica DMi8 S with an HCX PL APO 63x/1.40 (Oil) objective. For MTT assays (*90*) 6000 cells of each condition were seeded per well in a 96-well plate in four replicates, incubated for 4 h at 37 °C after the addition of 1 mM MTT (3-(4,5-Dimethylthiazol-2-yl)-2,5-Diphenyltetrazolium Bromide, ThermoFisher, M6494), treated with DMSO for 10 min at 37 °C and finally their absorbance was measured at 540 nM in an Infinite® 200 PRO plate reader (Tecan). For RNA-sequencing, one million cells of each condition were lysed in Trizol (ThermoFisher, 15596018) and RNA was isolated with the Direct-Zol RNA mini-prep kit (ZymoResearch, R2052). Elongation rates for wild-type and mutant samples were calculated as described in the section Transcriptional elongation speed.

### Eukaryotic Cell Lines

Human umbilical vein endothelial cells (HUVECs) from individual healthy donors were purchased by Lonza Inc.; human primary lung fibroblasts (IMR90) from two different isolates were obtained via the Coriell repository. All these lines were biannually checked for mycoplasma contamination and tested negative.

### Animal Strains Used & Animal Ethics

*Mus musculus*. Female F1 hybrid mice (C3B6F1) were generated in-house by crossing C3H/HeOuJ females with C57BL/6 NCrl males (strain codes 626 and 027, respectively, Charles River Laboratories). The DR study involving live mice was performed in accordance with the recommendations and guideline of the Federation of the European Laboratory Animal Science Association (FELASA, EU directive 86/609/EEC), with all protocols approved by the Landesamt für Natur, Umwelt und Verbraucherschutz, Nordrhein-Westfalen (LANUV), Germany (reference numbers: 8.87-50.10.37.09.176, 84-02.04.2015.A437, and 84-02.04.2013.A158) and the Netherlands (IACUC in Bilthoven, NIH/NIA 1PO1 AG 17242).

*Drosophila melanogaster*: v[1], *RpII215*[4] (RRID:BDSC_3663) mutant flies and *Repo-Gal4*(*91*) were obtained from the Bloomington Drosophila Stock Center (NIH P40OD018537). The *RpII215*[4] allele and *Repo-Gal4*(*91*) were backcrossed for 6 generations into the outbred *white Dahomey* (*wDah*) wild type background (Grönke et al., 2010) generating the *wDah, RpII215*[4] stock, which was used for experiments. *wDah, dilp2-3,5* flies (RRID:BDSC_30889) (Grönke et al., 2010) and *UAS-Histone 3* (*UAS-H3*) were previously generated in the lab and backcrossed for 6 generations into the outbred *wDah* wild type background. Female flies were used for all experiments.

*C. elegans* strains used: AA4274 ama-1(m322), ama-1(syb2315), CB1370 daf-2(e1370), N2 wild type.

### Human Research Participants & Human Ethics

Participants were searched using bulletins in which healthy individuals interested in taking part in a study examining aging-related changes were asked to contact the Dept. 2 of Internal Medicine (UoC) by telephone. A trained employee ruled out relevant pre-existing diseases using a structured questionnaire. The test persons were then invited for an appointment at the University Hospital Cologne to obtain the blood samples used for sequencing in the study at hand. Ethics were reviewed by the Institutional Review Board, Medical Faculty, University of Cologne. Permission was granted on Jan-17, 2018, Proposal ID: 17-362. The study was registered to the “Deutsches Register Klinischer Studien (DRKS)” (DRKS00014637).

## Code Availability

Standard procedures and published codes were used for data analysis (see **Methods** above). Custom code created for processing the data is available from https://github.com/beyergroup/ElongationRate.

## Figures

Species silhouette images were downloaded from phylopic.org. Figures were generated using Adobe Illustrator and Inkscape.

## Data availability

RNA-seq data used in this study is available at GEO database under accession numbers GSE102537 and GSE92486.

## Acknowledgements

We acknowledge the Bloomington Drosophila Stock Center for fly stocks, and the Max-Planck-Genome-Centre Cologne and the Cologne Center for Genomics for library preparation and sequencing. We thank Martijn Dollé for providing mice used in the aging kidney experiment. We thank Hans Schöler for sharing cell lines. We acknowledge funding from the Max Planck Society (to LP and AA), Bundesministerium für Bildung und Forschung Grant SyBACol 0315893A-B (to TB, AB and LP) and the European Research Council under the European Union’s Seventh Framework Programme (FP7/2007-2013)/ERC grant agreement number 268739 to LP. We also acknowledge support by the Deutsche Forschungsgemeinschaft (DFG) via the SPP1935 (Project 313408820) and Basic module grants (290613333 and 285697699) to AP.

## Corresponding authors

Correspondence to: Adam Antebi, Argyris Papantonis, Linda Partridge, and Andreas Beyer.

