## Supplementary tables and figures for "Aging-associated changes in transcriptional elongation influence metazoan longevity"

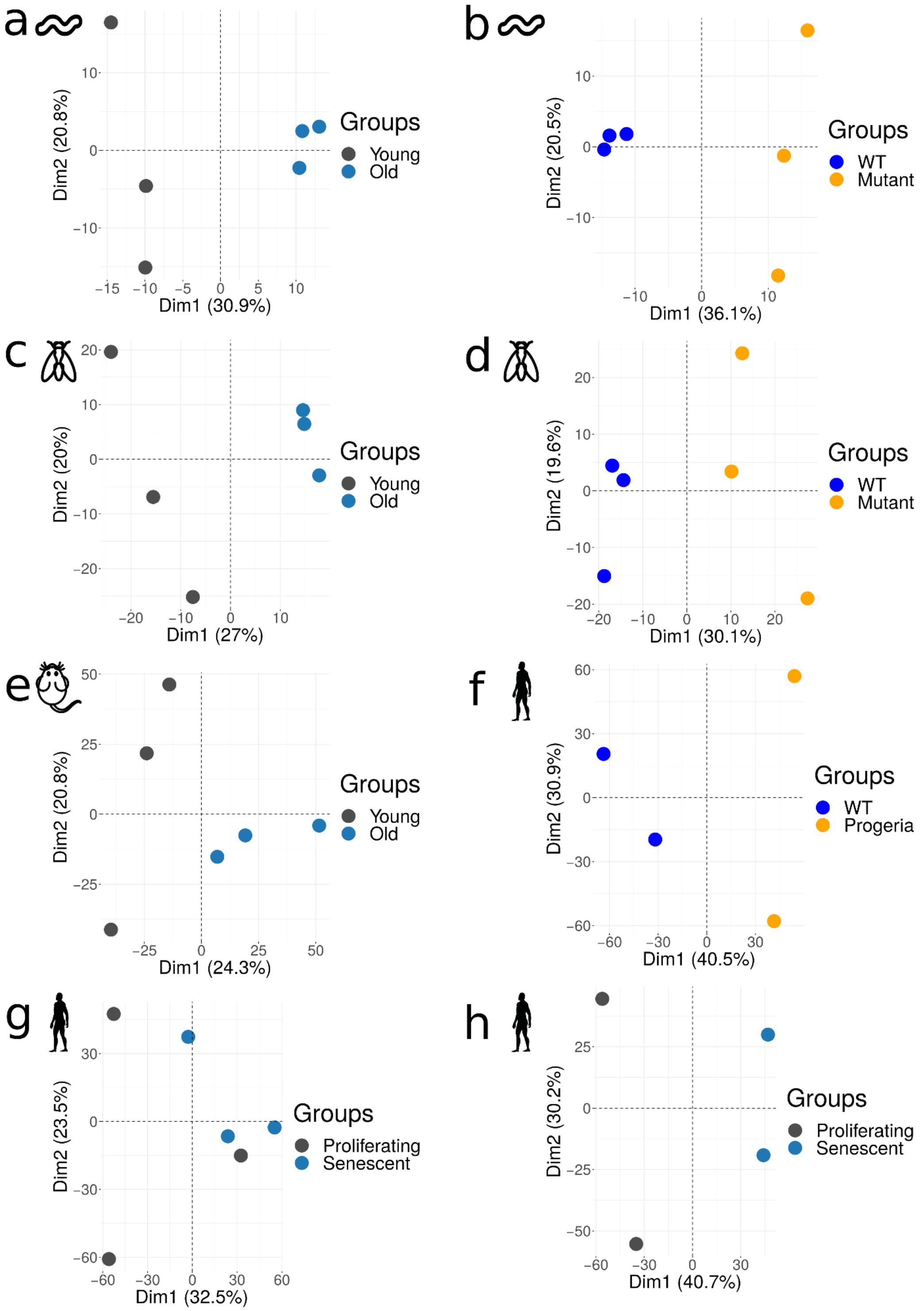


**Extended Data Figure 1: PCAs of slopes of intronic read distribution.** Principal component analysis (PCA) of the slopes of *C. elegans* (**(a)** wt 21 d vs 1 d; **(b)** 14 *ama-1(m322)* d vs wt 14 d), *D. melanogaster* (**(c)** wt heads 50 d vs 10 d, **(d)** RpII215[4] heads 50 d vs wt 50 d), *M. musculus* (**(e)** kidney: 24 mo vs 3 mo), *H. sapiens* (**(f)** Progeria: WT vs progeria mutants, **(g)** HUVEC and **(h)** IMR90: Senescent vs Proliferating).

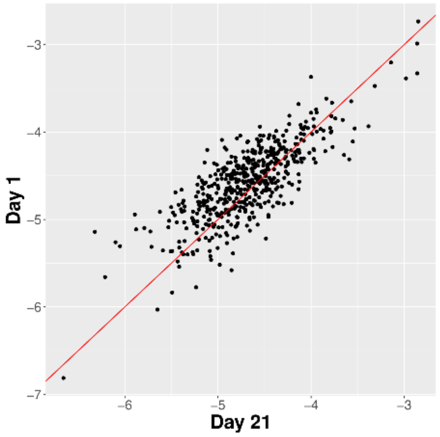
Day 1
Day 21

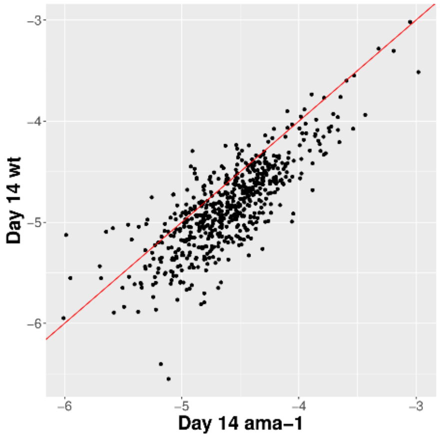
Day 14 wt
Day 14 ama−1

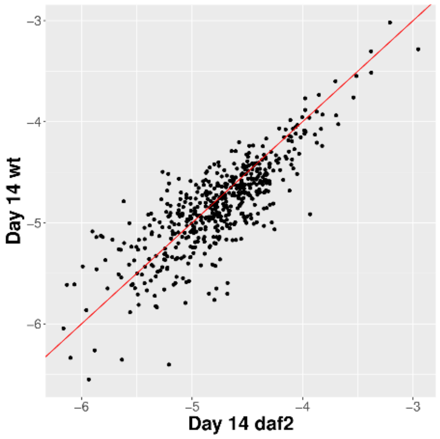
Day 14 wt
Day 14 daf2

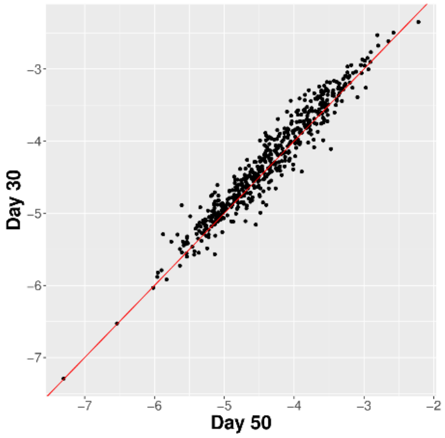
Day 30
Day 50

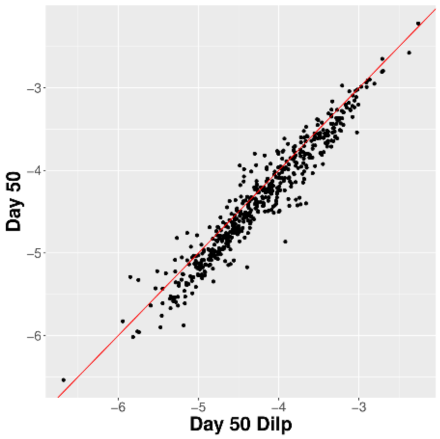
Day 50
Day 50 Dilp

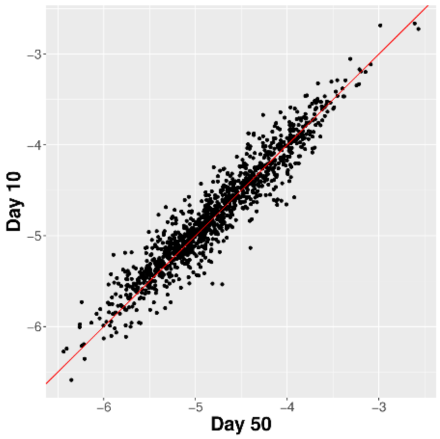
Day 10
Day 50

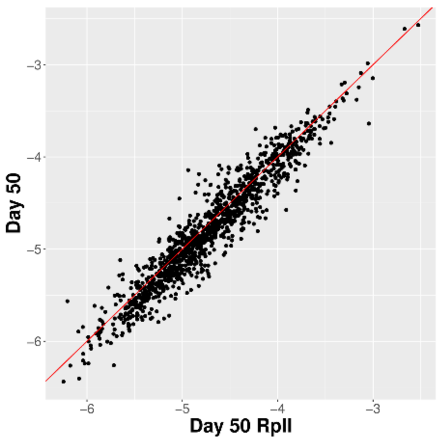
Day 50
Day 50 RpII

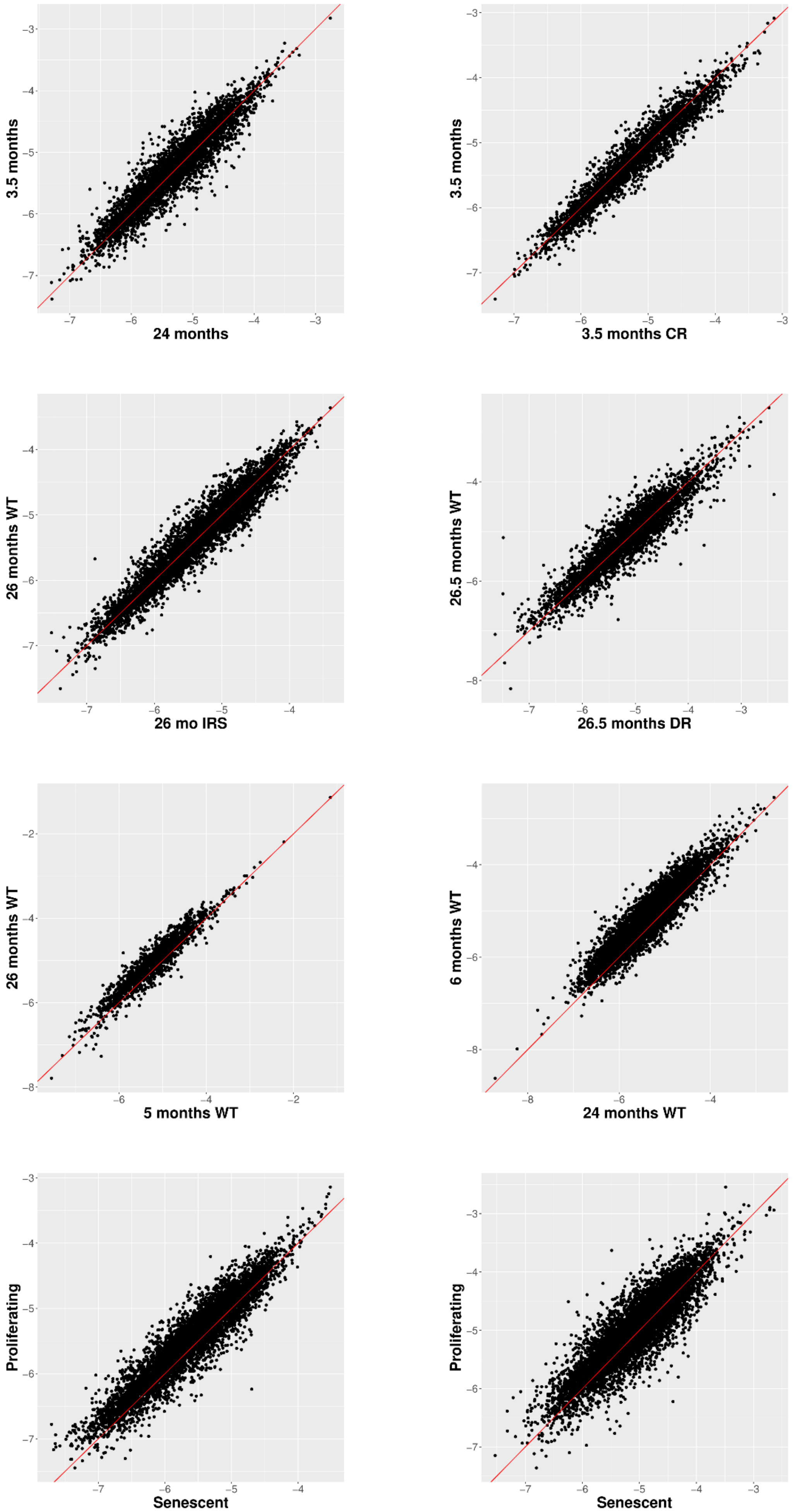
3.5 months
24 months
3.5 months
3.5 months CR
26 months WT
26 mo IRS
26.5 months WT
26.5 months DR
26 months WT
5 months WT
6 months WT
24 months WT
Proliferating
Senescent
Proliferating
Senescent

**Extended Data Figure 2:** Scatterplots of intronic slope(-log10) for each condition and species **(C. elegans, D. melanogaster, M.musculus, R. norvegicus, H. sapiens).**

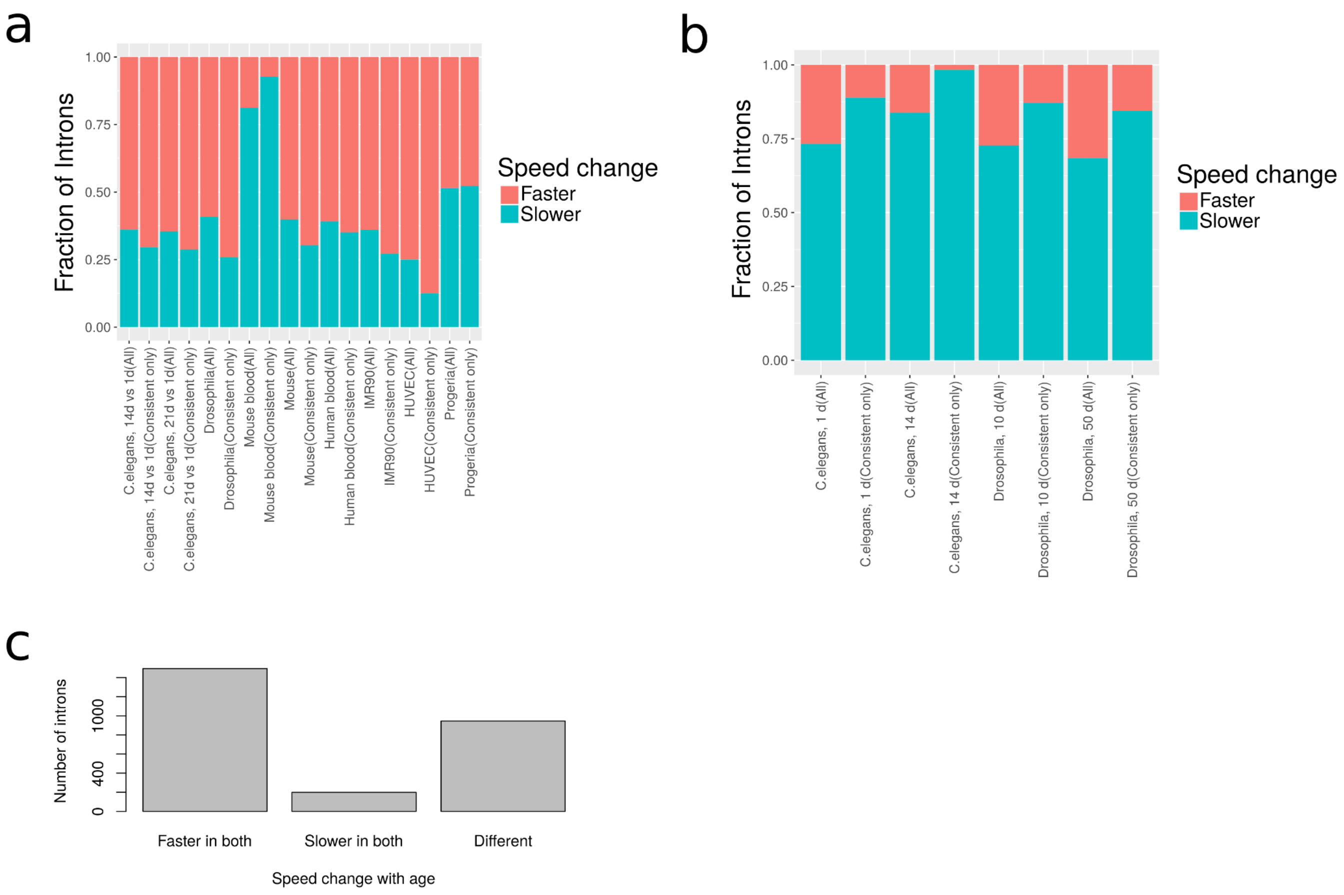


**Extended Data Figure 3: Consistency of RNA Pol-II speed changes. (a)** Change of elongation rate with aging or senescence in introns of *C. elegans*, *D. melanogaster*, *M. musculus* and *H. sapiens,* before and after filtering for introns that consistently change in speed in all replicates.**(b)** Change of elongation rate with mutations that slow down the speed of RNA-Pol-II in introns of *C. elegans* and *D. melanogaster* before and after filtering for introns that consistently change in speed in all replicates.**(c)** Comparison of the change of elongation rate  with aging between IMR90 and HUVEC using the same introns.

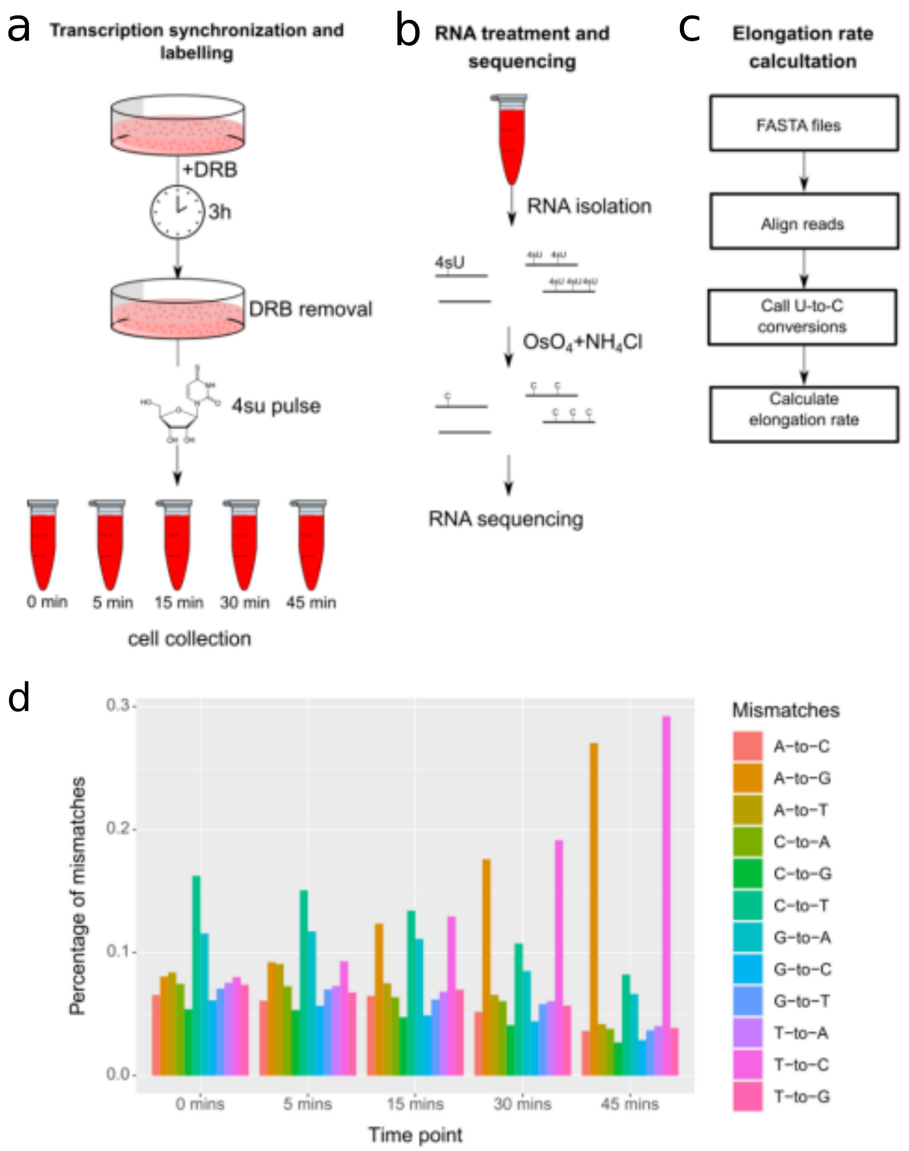


**Extended Data Figure 4: 4SU-DRB labelling and TUC conversion to calculate RNA-Pol-II elongation rate. (a-c)** Schematic representation of the 4SU-DRB labelling (**a**), TUC conversion (**b**) and elongation rate calculation (**c**). **(d)** Percentage of mismatches in every time point of the experiment (0 mins, 15 mins, 30 mins, 45 mins) in one of the proliferating replicates. There is a noticeable increase in A-to-G and T-to-C mismatches in the last two time points.

a

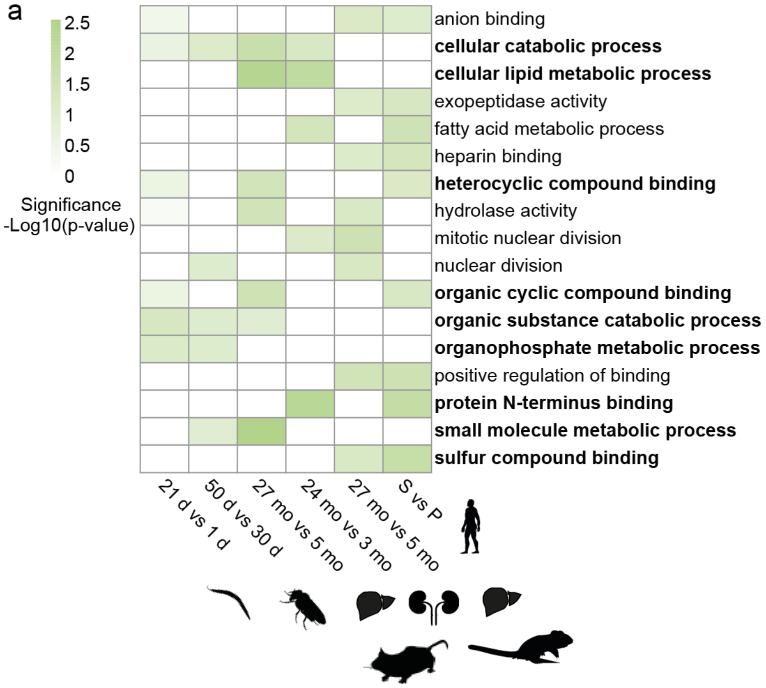


b

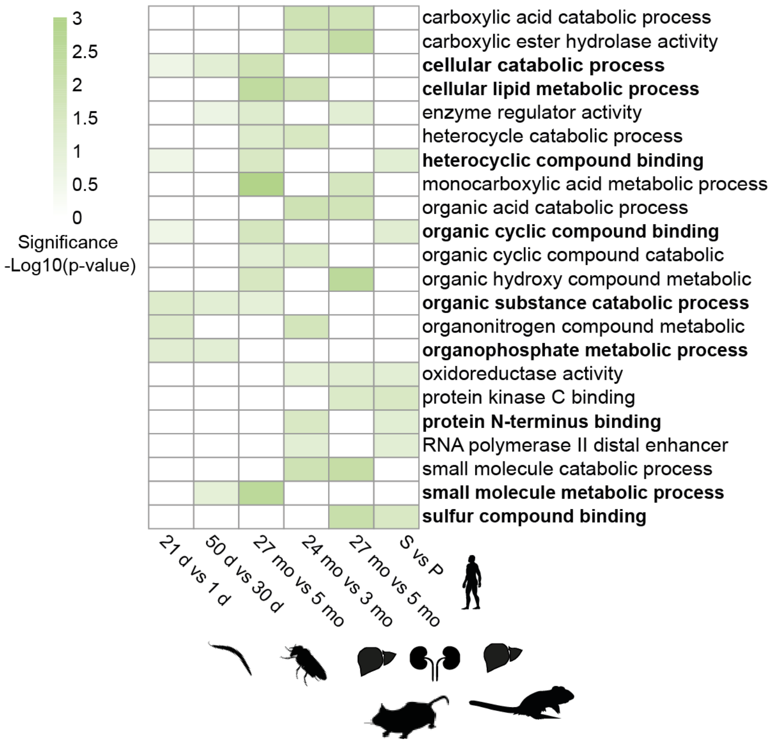


**Extended Data Figure 5: Genes with increase in Pol-II speed are associated with metabolism and catabolism related pathways.** GO enrichment analysis of genes with increased Pol-II speed across species: *C. elegans* (21 d vs 1 d)*, D. melanogaster* (heads: 50 d vs 30 d)*, M. musculus* (kidney: 24 mo vs 3 mo)*, R. norvegicus* (liver: 24 mo vs 6 mo), *H. sapiens* (IMR90: Senescent vs Proliferating). GO enrichment of (**a**), top 200 (**b**), top 300 genes with an increase in Pol-II speed change for each species (common terms between the two sets in bold). Color scale indicates the significance of the enrichment (all GO terms enriched with p-values below 0.05, with at least 10 significant genes for each GO categories, Fisher elim test).

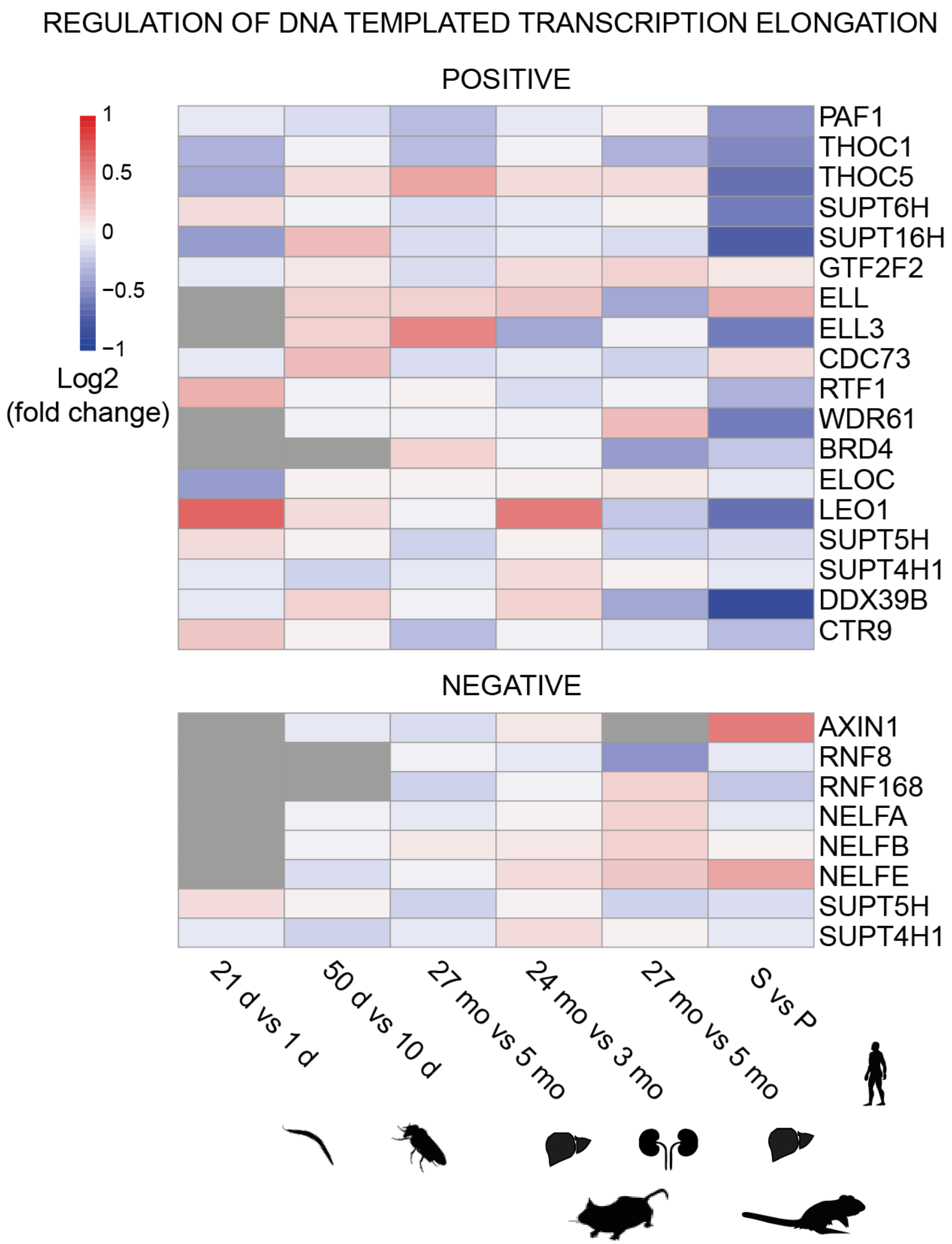


**Extended Data Figure 6: Heatmap of differential expression (log$_2$ fold change) of MSigDB** *(61)* **annotated genes for 'regulation of DNA templated transcriptional elongation'**. **Top**: activators of transcriptional elongation (POSITIVE); **Bottom**: repressors of transcriptional elongation (NEGATIVE). Data shown for *WT* aging time courses: worm (21 d vs 1 d), fly heads (50 d vs 10 d), mouse liver (27 mo vs 5 mo), mouse kidneys (24 mo vs 3 mo) and human fibroblast cell line (IMR90: Senescent vs proliferating).

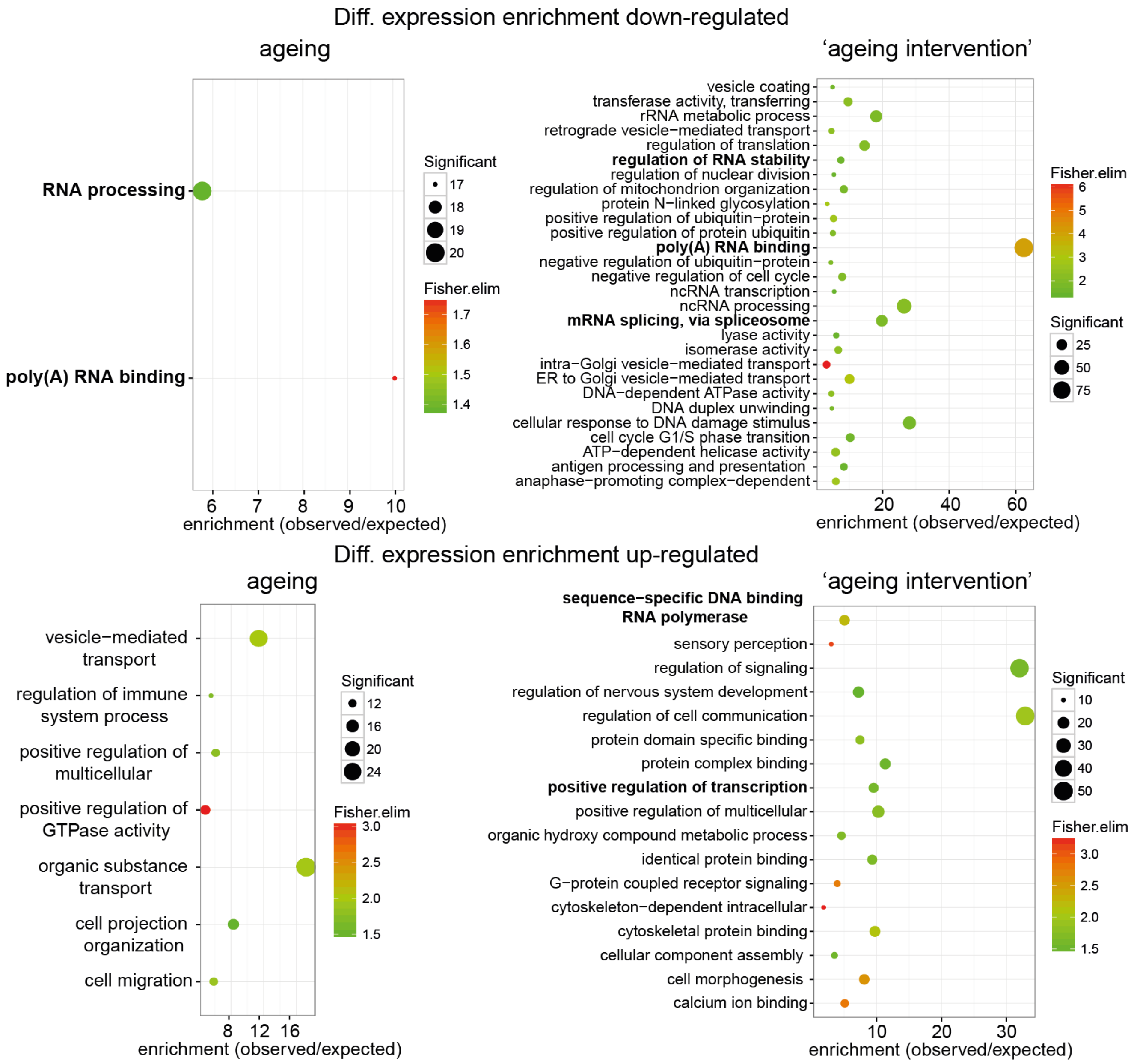


**Extended Data Figure 7: Functional enrichment for across-species differential expression analysis.** GO enrichment for consistently down-regulated (top) or up-regulated (bottom) genes across species during aging (left) or 'aging intervention' (right) (aging up-regulated: 92 genes; aging down-regulated: 71 genes; 'aging intervention' up-regulated: 164 genes; 'aging intervention' down-regulated: 473 genes; as background for the enrichment analysis a set of 4784 orthologue genes between *H. sapiens*, *R. norvegicus*, *M. musculus*, *D. melanogaster*, *C.* elegans was used. All p-values $*P < 0.05$, significant genes > 10, fisher elim test). GO terms related to transcription and splicing are indicated in bold.

a

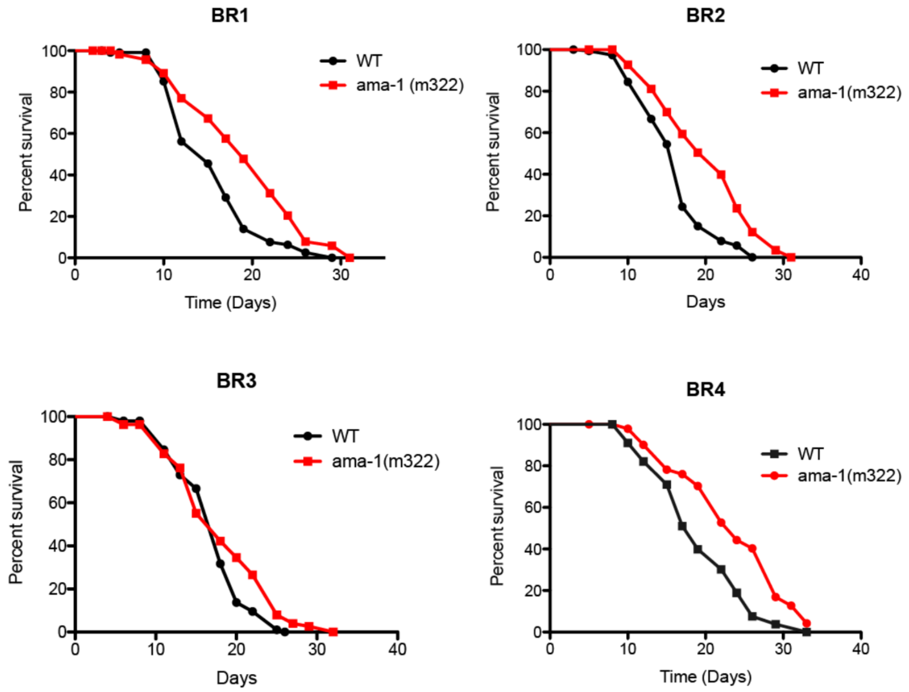


b

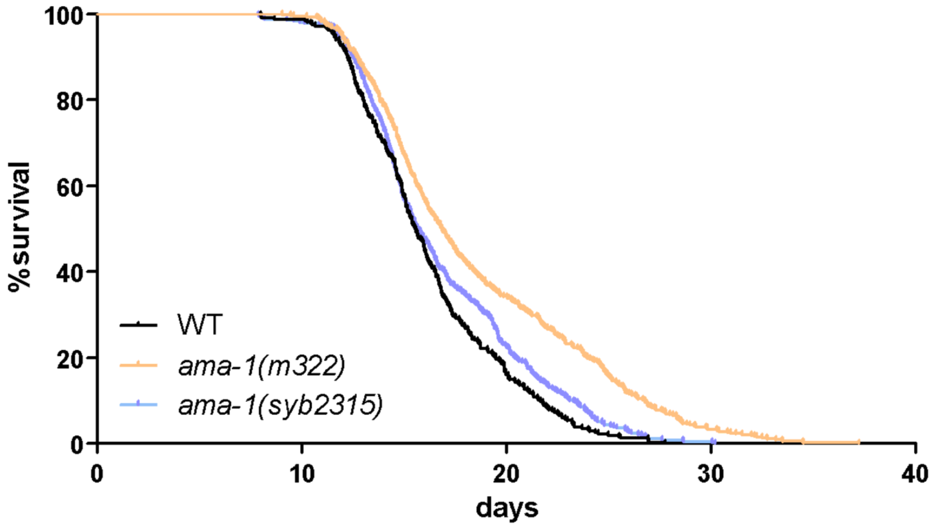


**Extended Data Figure 8: Slowing down Pol-II in *C. elegans* increases lifespan**. **(a)** Survival of wild-type and *ama-1(m322)* mutant worms conferring a slow Pol-II elongation rate (4 replicates, BR1:1.267, P < 0,0001; BR2:1.23,P < 0.0001; BR3:1, P = 0.0342; BR4:1.263,P < 0.0001, log-rank test, Mantel-cox). **(b)** C. elegans lifespan analysis after CRISPR/Cas9 mediated reversion of the slow RNAPII mutation. Survival curves of the strain harbouring the slow RNAPII mutation (ama-1 m322) and wild-type controls compared to worms after CRISPR/Cas9 engineered reversion of the slow mutation back to the wild type allele (ama-1 syb2315). Animals with slow Pol-II have a significantly increased lifespan. CRISPR/Cas9 engineered reversion restored lifespan essentially back to wild-type levels. (3 replicates; n > 300 per strain).

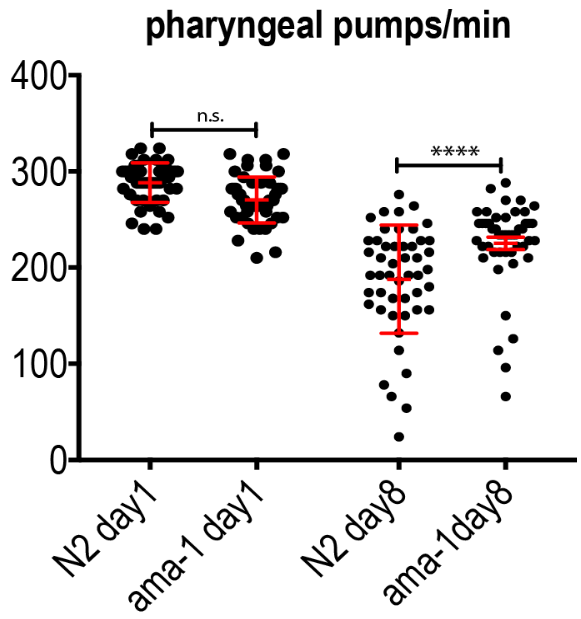


**Extended Data Figure 9: Slowing down Pol-II in C. elegans ameliorates the age-related decline in pharyngeal pumping rates.** Pumping rates of wild type N2 and ama-1 mutant worms were measured on day 1 and day 8. Pumping rates were not significantly different on day 1, but ama-1 worms showed higher pumping rates compared to wild types on day 8, suggesting that the mutant worms are healthier at old age.

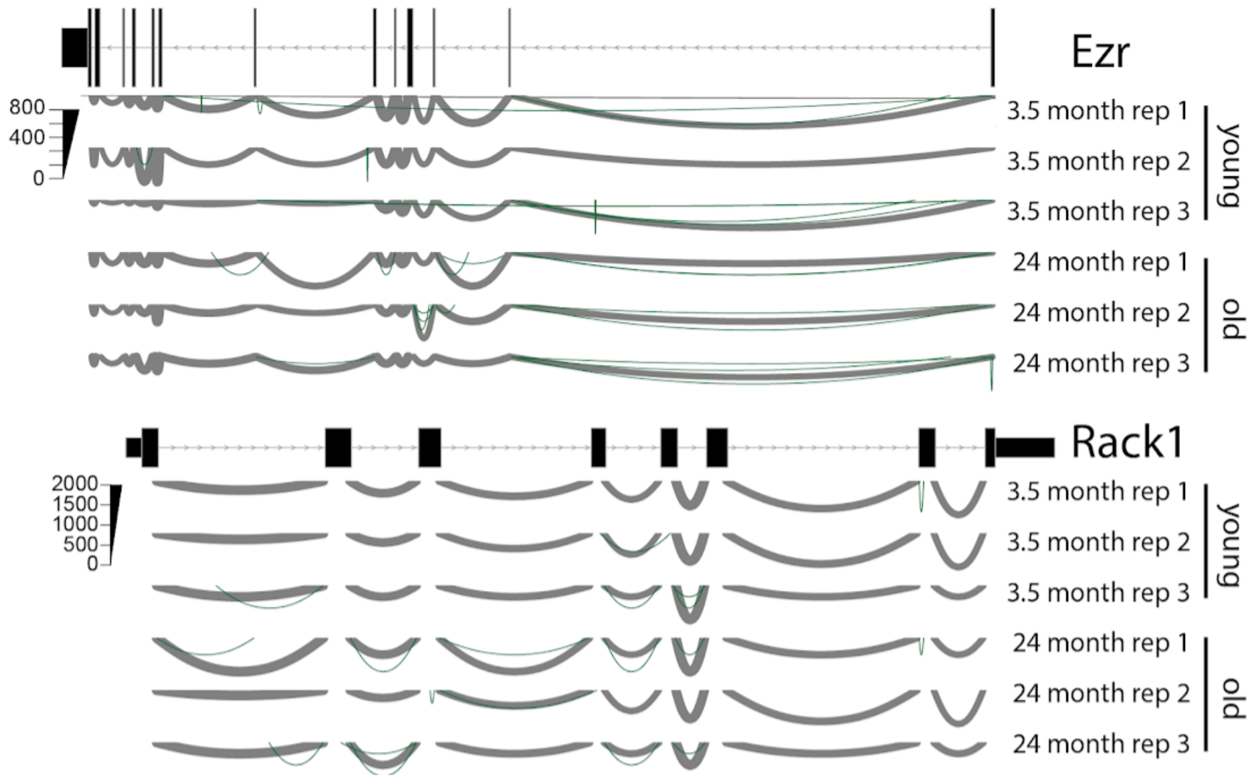

**Extended Data Figure 10: Examples of rare splice site changes** for gene Ezr and Rack1 with 3 replicates young (3.5 month) and old (26 month). Line thickness encodes the number of reads supporting this junction. Rare splice sites are shown in green**.**

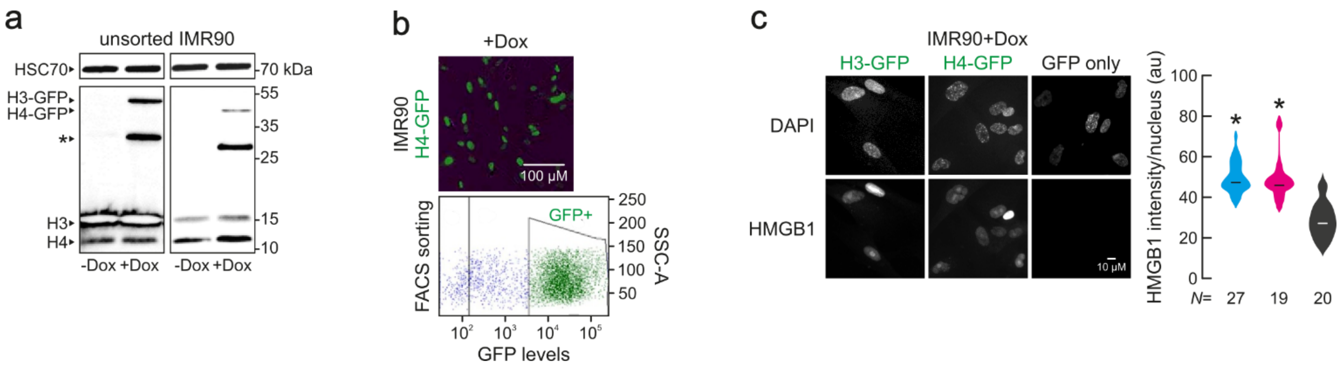


**Extended Data Figure 11: H3-GFP and H4-GFP overexpression in IMR90 cells. (a)** Western blot experiments confirm the overexpression of the H3-GFP and H4-GFP proteins. **(b)**Visual confirmation of the Dox induction of H3/H4 expression and FACS sorting of GFP-positive cells. **(c)** Typical immunofluorescence images of H3-GFP, H4-GFP and control IMR90 cells(*left)* show increased DAPI levels in histone overexpression nuclei. Violin plots(*right*) quantify this reduction. N specifies the number of cells analyzed per condition.

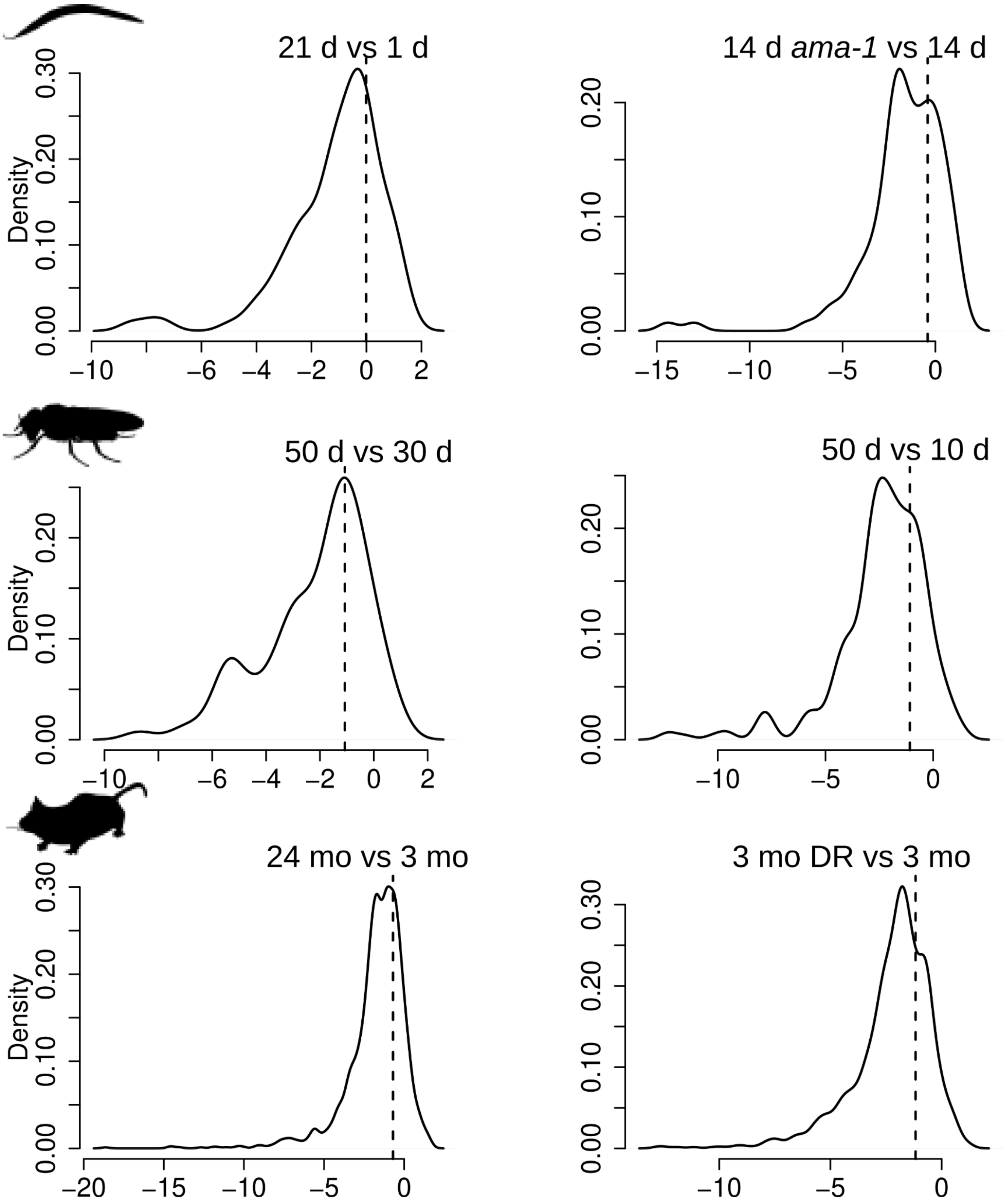

21 d vs 1 d
Density
0.00
0.10
0.20
0.30
−10
−6
−4
−2
0
2
14 d *ama-1* vs 14 d
0.00
0.10
0.20
−15
−10
−5
0
50 d vs 30 d
Density
0.00
0.10
0.20
−10
−6
−4
−2
0
2
50 d vs 10 d
0.00
0.10
0.20
−10
−5
0
24 mo vs 3 mo
Density
0.00
0.10
0.20
0.30
−20
−15
−10
−5
0
3 mo DR vs 3 mo
0.00
0.10
0.20
0.30
−10
−5
0

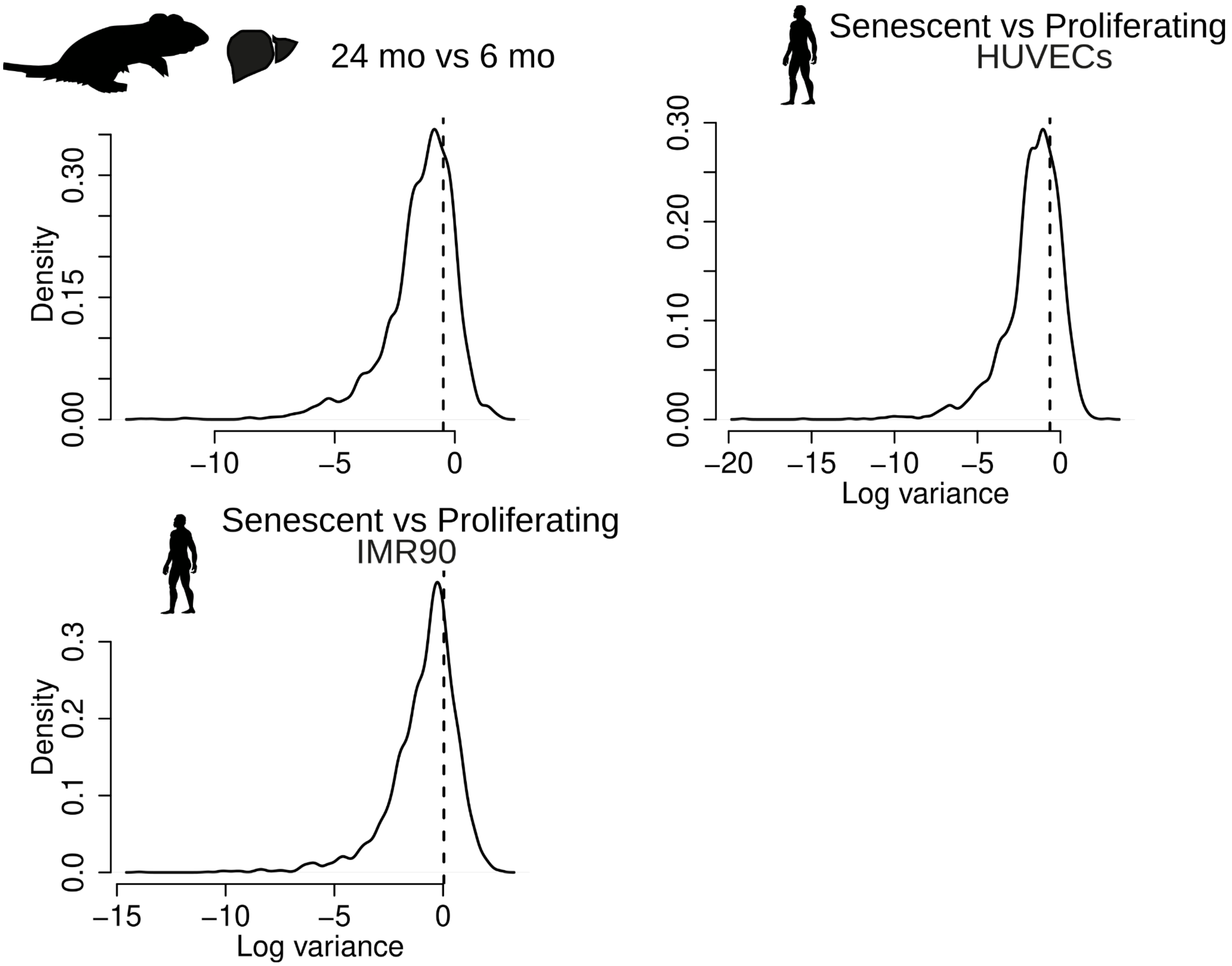


**Extended Data Figure 12**: **Variation of Pol-II elongation speed changes for different introns of the same gene.** Distribution of variances of Pol-II speed estimates (slope per intron) for introns within the same gene. Average variance of speed estimates across all introns (i.e. between genes; global average) is shown as a dashed vertical line for *C. elegans* (21 d vs 1 d; 14 *daf-2* d vs 14 d)*, D. melanogaster* (heads 50 d vs 30 d; 50 d vs 10 d)*, M. musculus* (kidney: 24 mo vs 3 mo; 3 DR mo vs 3 mo)*, R. norvegicus* (liver: 24 mo vs 6 mo), *H. sapiens* (Umbilical vein endothelial (HUVECs); fibroblast fetal lung (IMR90): Senescent vs Proliferating). The vast majority of intra-gene variances are below the average inter-gene variance, suggesting that introns of the same gene have coupled Pol-II elongation speeds.

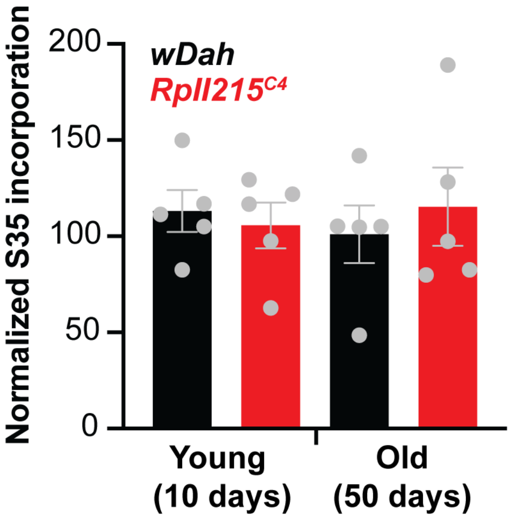


**Extended Data Figure 13: Protein biosynthesis rates do not change with aging in Drosophila:** Ex-vivo S35 incorporation assay shows no significant difference in translation rates in female fly heads between wDah control and RpII215C4 mutants both at young (10days) and old age (50 days). N=5 biological replicates with 25 heads per replicate.

**Table 1**

| Species | Comparison | Distance from promoter | Intron length | Gene expression log2FC | Circular RNA index |
|---|---|---|---|---|---|
| *C. elegans* | 21 d vs 1 d | 0.014 | 0.071 | -0.266 | -0.085 |
| *C. elegans* | 14 d ama-1 vs 14 d wt | 0.008 | -0.110 | -0.247 | 0.160 |
| *C. elegans* | 14 d daf2 vs 14 d wt | 0.020 | 0.007 | -0.278 | 0.130 |
| *D. melanogaster* | 50 d vs 10 d | -0.036 | 0.019 | -0.023 | 0.019 |
| *D. melanogaster* | 50 d RpII215 vs 50 d | 0.002 | -0.091 | -0.161 | -0.043 |
| *D. melanogaster* | 50 d dilp 2,3-5 vs 50 d | -0.133 | -0.170 | 0.041 | -0.092 |
| *M. musculus* | 24 mo vs 3.5 mo | 0.011 | 0.043 | -0.230 | -0.045 |
| *H. sapiens* | Senescent vs Proliferating (IMR90) | -0.021 | 0.046 | -0.274 | 0.184 |
| *H. sapiens* | Senescent vs Proliferating (HUVEC) | 0.014 | 0.036 | -0.289 | 0.020 |

**Extended Data Table 1:** Table of correlations(Pearson correlation) between the change in elongation rate and characteristics of the introns in which the elongation rate was measured in selected RNA-SEQ datasets.

**Table 2**

| **Species** | **Tissue** | **Enrichment protocol** | **Sequencing parameters** Paired-/single-end, read length, millions of reads (M) | **Time points[1] and conditions** |
|---|---|---|---|---|
| *C. elegans* | Whole body | TruSeq Stranded Total RNA Library | Paired-end, 75 *bp*, 25 M | Day 1 (*WT*), day 7 (*WT*), day 14 (*WT*), day 21 (*WT*) |
| | Whole body | TruSeq Stranded Total RNA Library | Paired-end, 75 *bp*, 25 M | Day 14 (*WT*, *daf-2(e1370), ama-1(m322)*) |
| | Whole Body | TruSeq Stranded Total RNA library | Paired-end, 75 bp, 25 M | Day 1 (WT, ama-1(m322)) |
| *D. melanogaster* | Head | TruSeq Stranded Total RNA Library | Single-end, 100 *bp*, 37.5 M | Day 30 (*WT*, *dilp2,3-5*), Day 50 (*WT*, *dilp2,3-5*) |
| | Head | TruSeq Stranded Total RNA Library | Paired-end, 75 *bp*, 30 M | Day 10 (*WT*, $RpII215^{C4}$), day 50 (*WT*, $RpII215^{C4}$) |
| *M. musculus* | Kidney | TruSeq Stranded Total RNA Library | Paired-end, 75 *bp*, 70 M | Month 3 (*WT*), month 24 (*WT*) |
| | Kidney | TruSeq Stranded Total RNA Library | Paired-end, 75 *bp*, 30 M | Month 3 (*WT*, *DR*) (4 replicates) |
| | Liver | TruSeq Stranded Total RNA Library | Paired-end, 75 *bp*, 37.5 M | Month 5 (*WT*, *DR*) |
| | Liver | TruSeq Stranded | Paired-end, 75 *bp*, 37.5 M | Month 16 (*WT*, *DR*) |

[1] Triplicate except where mentioned otherwise.

| | | | | |
|---|---|---|---|---|
| | | Total RNA Library | | |
| | Liver | TruSeq Stranded Total RNA Library | Paired-end, 75 *bp*, 37.5 M | Month 27 (*WT*, *DR*) |
| | Blood | TruSeq Stranded Total RNA Library | Paired-end, 75 *bp*, 70 M | Month 5 *(WT),* month 27 *(WT)* |
| | Hypothalamus | TruSeq Stranded Total RNA Library | Single-end, 100 *bp*, 30 M | Month 26 (*WT*, *IRS1*-/-) |
| *R. norvegicus* (*17*) | Liver | TruSeq Stranded Total RNA Library | Single-end, 50 *bp*, 60 M | Month 6 (*WT*), month 24 (*WT*) (2 replicates) |
| | Brain[2] | TruSeq Stranded Total RNA Library | Single-end, 50 *bp*, 20 M | Month 6 (*WT*), month 24 (*WT*) (6 replicates) |
| *H. sapiens* | Fetal lungs (IMR90) | Nascent RNA | Paired-end, 75 *bp*, 25 M | Early passage, late passage (2 replicates) |
| | Fetal lungs (IMR90) | TruSeq Stranded Total RNA Library | Paired-end, 75 *bp*, 50 M | Early passage, late passage (2 replicates) |
| | Umbilical vein endothelial (HUVECs) | Nascent RNA | Paired-end, 75 *bp*, 50 M | Early passage, late passage (2 replicates) |
| | Umbilical vein endothelial (HUVECs) | TruSeq Stranded Total RNA Library | Paired-end, 75 *bp*, 100 M | Early passage, late passage |
| | Fibroblast skin | TruSeq Stranded Total RNA Library | Paired-end, 100 *bp*, 100 M | Progeria patient (HSS) (2 replicates) |

[2] Not included in the analysis due to low coverage (bellow 1X genome coverage or 29 M sequenced reads for *R. norvegicus* ; genome coverage calculated using Lander-Waterman formula)

| | | | | |
|---|---|---|---|---|
| | | | Paired-end, 75 *bp*, 100 M | Healthy donor, sex/age matched with progeria patient (2 replicates) |
| | Blood | TruSeq Stranded Total RNA Library | Paired-end, 75 *bp*, 70 M | Healthy donor, 6 females, 6 males, age range: 21-70 |

**Extended Data Table 2:** Description of the RNA-seq datasets used in the study.
